# Spermidine alleviates depression via control of the stress response

**DOI:** 10.1101/2024.12.04.626625

**Authors:** Sarah Mackert, Christine Niemeyer, Yara Mecdad, Tim Ebert, Thomas Bajaj, Sylvère Durand, Sebastian J. Hofer, Andreas Zellner, Bianca Besteher, Jan Engelmann, Barbara Gisabella, Valerie Kempf, Lily A. Heckmann, Marcel Laakmann, Emily L. Newman, Clara Sokn, Daniel E. Heinz, Ellen Junglas, Andrés Uribe-Marino, Christoph Magnes, Claudia Klengel, Anna Müller, Lydia Opriessnig, Andreas Meinitzer, Martina Lennarz, Katja M. Wirtz Martin, Rawan Albatarni, Marieke Brockherde, Klaus Lieb, Henrik Rohner, Birgit Stoffel-Wagner, Alexandra Philipsen, Bernhard Kuster, Markus Kölle, Kerry J. Ressler, Nils Opel, Mathias V. Schmidt, Harry Pantazopoulos, Marianne B. Müller, Guido Kroemer, Tobias Eisenberg, Jakob Hartmann, Frank Madeo, Nils C. Gassen

## Abstract

Depression is a stress-associated disorder, and it represents a major global health issue. Its pathophysiology is complex and remains insufficiently understood, with current medications often showing limited efficacy and undesirable side effects. Here, we identify imbalanced polyamine levels and dysregulated autophagy as key components of the acute stress response in humans, and as hallmarks of chronic stress and depressive disorders. Moreover, conventional antidepressant pharmacotherapy increases endogenous plasma concentrations of the polyamine spermidine exclusively in patients who respond to the treatment, suggesting a link between spermidine and successful outcomes. In a clinical trial, involving drug-naive depressed individuals, three weeks of spermidine supplementation increased autophagy and alleviated symptoms of depression. Behavioral and mechanistic findings of spermidine supplementation were validated in various mouse stress and depression models. In summary, spermidine supplementation mitigates polyamine dysregulation and stimulates autophagy under pathological stress conditions, offering a novel and well-tolerated treatment approach for stress-related depressive disorders.

## Introduction

Stress may be defined as a state of disrupted homeostasis triggered by an array of internal or external stressors, spanning physiological factors like inflammation, hypoxia, or heat shock, as well as psychological stressors such as interpersonal conflicts or traumatic experiences. The bodily reaction to a variety of stressors is characterized by the fine-tuning of intricate interactions between endocrine, metabolic, immune, neuronal, and behavioral systems, orchestrated by the hypothalamic-pituitary-adrenal axis and the sympathetic adrenal medullary axis^1^. These multi-layered adaptive responses are essential for overcoming acute stressful challenges, with the ultimate goal of restoring and maintaining organismal and cellular homeostasis. However, prolonged exposure to stressors impairs the cellular and systemic capacity to regulate these responses, leading to a chronic dysregulation of homeostatic mechanisms. Ultimately, an inadequate response to stress, resulting from the dysregulation of stress-responsive systems, contributes to the onset of severe and debilitating psychiatric conditions known as stress-related psychiatric disorders (SRPDs), with major depressive disorder (MDD) being the most common^2^.

Despite decades of research, mechanism-based pharmaceutical treatments for MDD are still rare, and currently available medications offer only limited effectiveness, with many patients failing to achieve full remission or experiencing a relapse after temporary improvement, while often causing undesirable side effects^3^. Yet, newly identified substances mostly lack discernible advantages over existing ones and frequently fail during the transition to clinical testing^4^. Moreover, drug discovery has predominantly focused on the central nervous system and neurotransmitter imbalances, neglecting the systemic nature of MDD and the various systemic and cellular regulatory levels that alter metabolism in crosstalk with the body and brain^5^. Autophagy is an intracellular catabolic process regulated by various factors, including nutritional cues such as amino acid metabolism, as well as energy demands and homeostatic needs^6^. It is functional in maintaining cellular homeostasis and is pivotal to the stress response pathways.

In this interdisciplinary, translational study, we first analyzed the mechanistic interplay between polyamine metabolism and autophagy in acute and chronic stress, using experimental models and clinical cohorts in mice and humans. Our initial findings from unbiased metabolomics revealed that acute stress causes an imbalance in polyamine metabolism and increases autophagic flux in both mice and humans. Additionally, we observed dysregulations in plasma and brain polyamine levels in subjects with depression compared to controls across three independent cohorts. Building on these findings, we conducted a double-blind, placebo-controlled study, in which spermidine (SPD) supplementation restored imbalances in polyamine metabolism and induced autophagic flux, significantly improving clinical symptoms. Our findings 1) identify the polyamine pathway as a key cellular hub modulating stress-related outcomes, 2) reveal novel mechanisms by which stressful life experiences are ultimately translated into alterations of the cellular metabolism, and 3) suggest SPD supplementation as a promising and well-tolerated treatment approach for depressive disorders that is low in side effects and cost-effective.

## Results

### Metabolic profiling uncovers activation of autophagy and modulation of polyamine metabolism in the acute human stress response

The acute stress response combines the complex interaction of many regulatory and molecular systems^7^, however, its precise metabolic fingerprint is not yet fully understood. We explored the metabolic response to acute stress by subjecting 22 healthy participants to a first-time 60-meter bungee jump (Supplementary Table 1, cohort #1)^8^. During the study, blood and saliva were obtained from participants at nine time points: one baseline sample seven days preceding the bungee jump, seven samples on the day of the bungee jump (two before and five after), and a final sample seven days after the bungee jump (Fig. 1A). A control group provided samples in a parallel setup without undergoing stress-exposure.

**Fig. 1.**
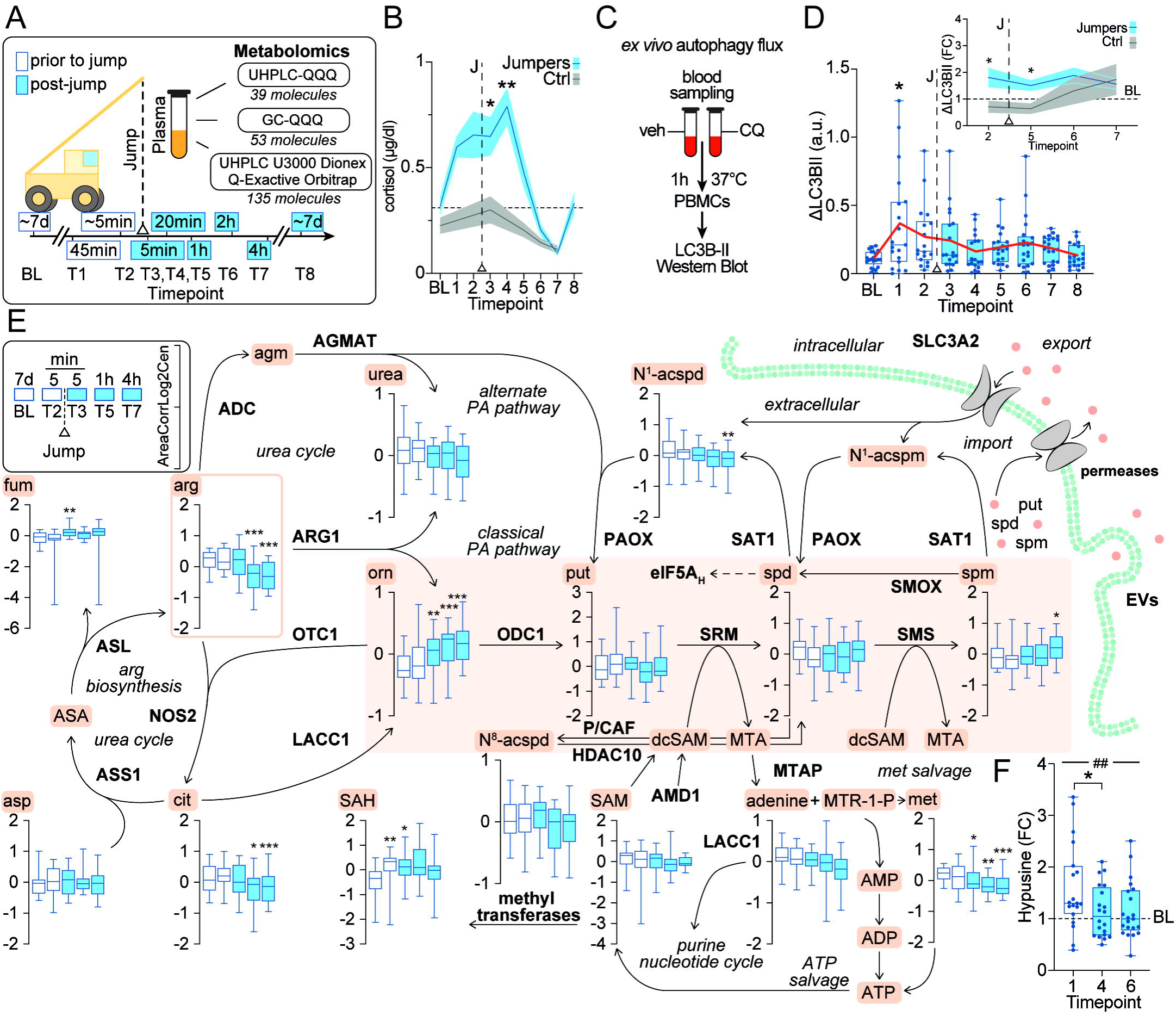
Acute stress induces autophagy and alters polyamine metabolism in humans. **(A)** Schematic overview of the study design for cohort #1, including all time points for blood sample collection and both unbiased and targeted metabolomic profiling of plasma samples using different analytical setups. **(B)** Absolute cortisol levels in saliva from bungee jumpers and control group throughout the study *(n = 22 Jumpers, n = 10 Ctrl; Sidak’s multiple comparisons test: T3: 95% CI: 0.03778 to 0.6579, p = 0.028; T5 = 95% CI: 0.06066 to 0.4830, p = 0.0067)*. **(C)** Schematic illustration of autophagic flux measurement in human blood samples. Whole blood samples were treated with either the lysosomal inhibitor chloroquine or a vehicle control and incubated for 1 hour at 37 °C. Afterwards PBMCs were isolated, and LC3B-II levels were measured by immunoblotting in the protein extracts from these cells. ΔLC3B-II indicates the difference in LC3B-II levels between chloroquine-treated samples and those treated with vehicle. **(D)** Autophagic flux represented as ΔLC3B-II from bungee jumpers and relative ΔLC3B-II levels in PBMCs from bungee jumpers and controls throughout the study *(n = 22 Jumpers; Sidak’s multiple comparisons test: BL vs. T1: 95% CI: −0.4074 to −0.004324, p = 0.043. Jumpers vs. Ctrl: n = 21 Jumpers, n = 10 Ctrl; Sidak’s multiple comparisons test: T2: 95% CI: 0.01092 to 0.9919, p = 0.0437; T5: 95% CI: 0.02764 to 0.9021, p = 0.0350)*. **(E)** Plasma levels of metabolites within the polyamine pathway and their associated enzymatic reactions in bungee jumpers at baseline, 5 minutes before the bungee jump, and 5 minutes, 1 hour, and 4 hours after the bungee jump *(A mixed effects model followed a two-stage linear step-up procedure of Benjamini, Kriger, and Yekutieli was used to correct for multiple comparisons by controlling the false discovery rate (< 0.05); comparisons to BL, n = 22. The box represents 25th to 75th percentiles, whiskers display the minimum to maximum values, and the line indicates median*. **(F)** Relative levels of hypusine normalized to vinculin in PBMCs from bungee jumpers before and after the bungee jump determined by immunoblotting *(Mixed effects model; main time effect: F_(2, 34)_ = 5.649, p = 0.0085. Tukey’s multiple comparisons test: T1 vs. T4: 95% CI: 0.08271 to 0.8345, p = 0.0160). Panel A-D and F: p ≤ 0.05 (*), p ≤ 0.01 (**), p ≤ 0.001 (***). Panel E: p ≤ 0.05 (*), p ≤ 0.01 (**), p ≤ 0.005 (***)).* **Abbreviations:** Abbreviations in alphabetical order: BL = Baseline, CI = confidence interval, Ctrl = control, # = Time effect. Abbreviations for analytical set ups: UHPLC-QQQ = Ultra-High-Performance Liquid Chromatography with Triple Quadrupole Mass Spectrometer, GC-QQQ = Gas Chromatography with Triple Quadrupole Mass Spectrometer. Abbreviations for autophagic flux measurements: LC3B-II = microtubule-associated protein 1 light chain 3 beta-II, PBMCs = peripheral blood mononuclear cells, CQ = chloroquine, veh = vehicle. Abbreviations for metabolic and regulatory enzymes and proteins: ADC = arginine deiminase complex, AMD1 = adenosylmethionine decarboxylase 1, ARG1 = arginase 1, ASL = argininosuccinate lyase, ASS1 = argininosuccinate synthase 1, EVs = extracellular vesicle s, GMAT = agmatinase, HDAC10 = histone deacetylase 10, LACC1 = laccase domain-containing 1, MTAP = methylthioadenosine phosphorylase, NOS2 = nitric oxide synthase 2, ODC1 = ornithine decarboxylase 1, OTC1 = ornithine transcarbamylase 1, PAOX = polyamine oxidase, P/CAF = p300/CBP-associated factor, SAT1 = spermidine/spermine N1-acetyltransferase 1, SLC3A2 = solute carrier family 3 member 2 (CD98), SMOX = spermine oxidase, SMS = spermine synthase, SRM = spermidine synthase. Abbreviations for metabolites and cofactors: ADP = adenosine diphosphate, agm = agmatine, AMP = adenosine monophosphate, arg = arginine, ASA = Argininosuccinic acid, Asp = Aspartate, ATP = adenosine triphosphate, Cit = citrulline, dcSAM = decarboxylated SAM, fum = fumarate, MTA = methylthioadenosine, MTR-1-P = methylthioribose-1-phosphate, N^1^-acspm = N1-acetylspermine, N^1^-acspd = N1-acetylspermidine, N^8^-acspd = N^8^-acetylspermidine, orn = ornithine, put = putrescine, SAH = S-adenosylhomocysteine, SAM = S-adenosyl-methionine, spd = spermidine, spm = spermine.

We observed an increase in salivary cortisol levels both in anticipation of the bungee jump and following the bungee jump (Fig. 1B). Cortisol levels returned to normal four hours after the bungee jump. To identify stress-responsive metabolic alterations in crosstalk with autophagic changes (Fig. 1C, D), we conducted unbiased metabolomic profiling using high-resolution mass spectrometry on plasma samples at five of the nine time points, which identified a total of 227 metabolic features selected for further analysis (Extended Data Fig. 1A, B, Supplementary Table 2). Notably, a major fraction of metabolites changed due to stress relative to the baseline measurement (Extended Data Fig. 1C). We further analyzed significantly altered metabolites to determine their association with specific regulatory pathways through Metabolite Set Enrichment Analysis (MSEA) based on the Kyoto Encyclopedia of Genes and Genomes (KEGG). Predominantly in the post-bungee jump phase, we observed a noticeable impact on metabolites related to the tricarboxylic acid cycle, and metabolites involved in the arginine (arg) biosynthetic pathways, which were distinctly affected one and four hours after the bungee jump, respectively (Extended Data Fig. 1D).

The amino acid arg serves as a precursor for polyamines, agmatine (agm), citrulline (cit) and urea (Fig. 1E). Post-bungee jump reductions in arg levels, accompanied by decreases in citrulline (cit), coincided with a significant rise in ornithine (orn) levels catalyzed by arginase 1 (ARG1), indicating a shift in arg metabolism towards the polyamine branch (Fig. 1E). Next, we investigated whether the stress-induced changes in the polyamine precursors arg and orn may affect the entire polyamine synthesis pathway, considering that the polyamine pathway contributes to cellular homeostasis through the induction of autophagy^9^. For this purpose, we conducted an additional targeted analysis of the three main polyamines, namely putrescine (put), spermidine (spd), and spermine (spm), as well as orn across all nine time points and in parallel to controls of the study, rather than just five as in the unbiased approach. This analysis confirmed the previously described stress-induced elevations in orn, while there were minor reductions in put and spd (Extended Data Fig. 1E).

Cellular metabolism - particularly of amino acids and polyamines - is tightly linked to the regulation of autophagy, and we and others have previously reported strong evidence for a direct impact of stress on autophagy in various models. To investigate whether this would also occur in humans, we next measured autophagic flux in peripheral blood mononuclear cells (PBMCs). For this purpose, we opted for an *ex vivo* approach in blood, which permits precise analysis without altering the metabolic state of the blood cells through cultivation (Fig. 1C)^10^. Immunoblotting of protein extracts from freshly isolated PBMCs revealed a significant induction of autophagy 45 minutes prior to the stress event, suggesting an effect of anticipated stress (Fig. 1D). In the post-stress phase, autophagy levels returned to baseline, paralleling the decline in cortisol levels (Fig. 1B, D, Extended Data Fig. 1F, G). Mechanistically, a hallmark of the link between polyamines and autophagy is increased hypusination. Consequently, in PBMC extracts, we observed an increase in hypusination levels before the bungee jump, suggesting a functional role for the polyamine-hypusination axis in translating psychological stress into cellular effects, such as autophagy modulation (Fig. 1F). The decrease in free spd in the plasma is thus likely due to its shuttling into the hypusine pathway (Extended Data Fig. 1E).

In conclusion, our human experiment, involving plasma and PBMC analyses, demonstrated that acute stress induces autophagic flux and modulates the polyamine-hypusination axis.

### Stress drives changes in polyamines and induces autophagy through glucocorticoid-receptor signaling

The metabolic alterations observed in the human acute stress model prompted further mechanistic inquiries into their underlying regulation. To address this and investigate whether the effects of acute stress on polyamine regulation are independent of the quality of the acute stressor, we extended our analyses to a single social defeat model (acute social defeat stress model, ASDS, mouse #1,^11^). Targeted metabolomics of polyamines and protein analyses in brain, liver, and plasma were performed following a single ASDS (Fig. 2A). In support of acute stress-induced autophagy, we demonstrated a degradation of sequestosome-1 (SQSTM1, best known as p62), which is an autophagic substrate and adaptor protein, in liver and brain tissue (Fig. 2B, Extended Data Fig. 2B). Consistent with autophagic flux data from the human trial, proteomic measurements of whole-brain extracts revealed a substantially increased proteome-wide fraction of down-regulated proteins in stressed mice, indicating elevated protein turnover likely mediated by autophagy (Fig. 2C, Extended Data Fig. 2A, Supplementary Table 3). Measurements of polyamines also aligned with the findings of the human study, showing a decline of spd and an increase in put (Fig. 2D, Extended Data Fig. 2C), while orn and spm remained unchanged (Extended Data Fig. 2D-F). On an enzymatic level, we show a significant induction of adenosylmethionine decarboxylase 1 (AMD1) in the liver and brain (Fig. 2E, Extended Data Fig. 2G), while the enzymes ODC1 and spermidine/spermine N1-acetyltransferase 1 (SAT1) remained unchanged (Extended Data Fig. 2H, I). Finally, and consistent with the effect observed after bungee jumping, we also detected increased hypusination levels in mice (Fig. 2F, Extended Data Fig. 2J). Thus both in mice and humans, acute stress leads to immediate channeling of the available cellular spd into hypusination, which causes autophagy.

**Fig. 2.**
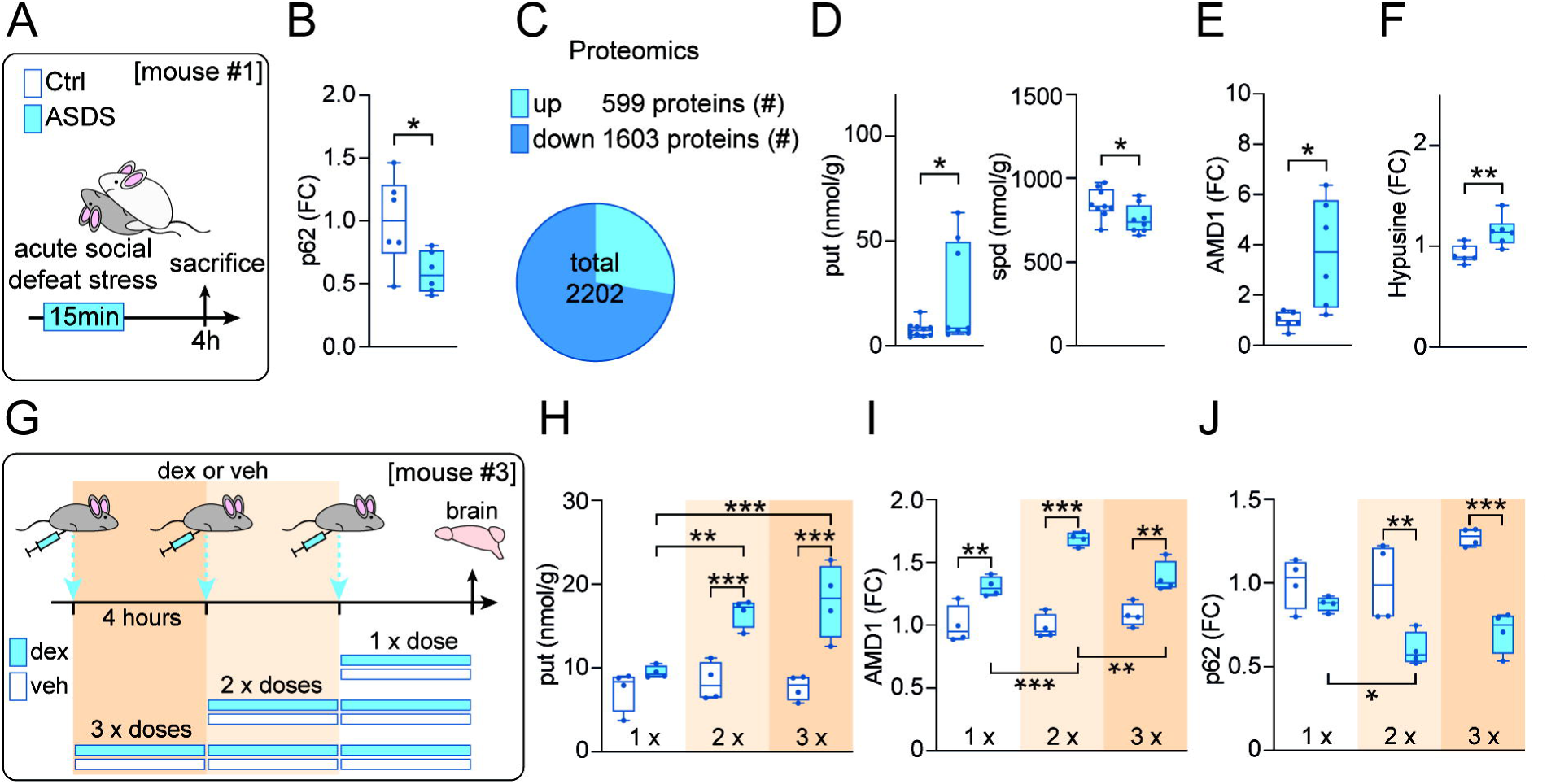
Glucocorticoid receptor signaling is involved in stress-induced changes in autophagy and polyamine metabolism. **(A)** Schematic overview of the acute social defeat stress (ASDS) paradigm in mice *(mouse #1: Western blots: n = 6 ASDS, n = 6 Ctrl; Polyamines: n = 11 ASDS, n = 11 Ctrl (Polyamine measurements included samples from mouse #2))*. **(B)** Relative p62 levels in liver lysates from mice in **(A)** determined by immunoblotting *(n = 6 ASDS, n = 6 Ctrl; unpaired two-tailed t-test: p = 0.0277)*. **(C)** Pie chart illustrating the differential expression of proteins in the brain in response to ASDS compared to controls. **(D)** Targeted analysis of put and spd levels in liver lysates from mice in **(A)** (put: n = 10 ASDS, n = 11 Ctrl; unpaired two-tailed t-test: p = 0.0292. spd: n = 8 ASDS, n = 9 Ctrl, unpaired two-tailed t-test: p = 0.0398). **(E)** Relative AMD1 levels in liver lysates from mice in **(A)** determined by immunoblotting *(n = 6 ASDS, n = 6 Ctrl; unpaired two-tailed t-test: p = 0.0125)*. **(F)** Relative levels of hypusine normalized to GAPDH in liver lysates from mice in **(A)** determined by immunoblotting *(n = 6 ASDS, n = 6 Ctrl; unpaired two-tailed t-test: p = 0.0096)*. **(G)** Schematic overview of the study design, including stepwise dex or veh injections administered intraperitoneally in mice *(mouse #3: n = 12 dex, n = 12 veh, with n = 4 receiving one dex injection, n = 4 receiving two dex injections, n = 4 receiving three dex injections, and n = 12 veh group was also divided into three groups and injected at the same intervals)*. **(H)** Targeted analysis of put levels in brain samples from mice in **(G)** (*n = 4 dex, n = 4 veh per injection; dex vs. veh: Sidak’s multiple comparisons tests: 3x: 95% CI: 5.797 to 14.95, p < .001; 2x: 95% CI: 3.672 to 12.83, p < .001. Tukey’s multiple comparisons tests: dex 2x vs. 1x: 95% CI: 2.684 to 11.57, p= 0.002; dex 3x vs. 1x: 95% CI: 4.109 to 12.99, p < 0.001).* **(I, J)** Relative AMD1 and p62 levels in brain lysates from mice in **(G)** determined by immunoblotting *(n = 4 dex, n = 4 veh. AMD1: dex vs. veh: Sidak’s multiple comparisons tests: 1x: 95% CI: 0,1135 to 0,4997, p = 0.002; 2x: 95% CI: 0,5100 to 0,8961, p = <.001, 3x: 95% CI: 0,1086 to 0,4947, p = 0.002. Tukey’s multiple comparisons tests: dex: 1x vs. 2x: 95% CI: −0,5688 to −0,1942, p = <. 001; 2x vs 3x: 95% CI: 0,1187 to 0,4932, p = 0.002. p62: dex vs. veh: Sidak’s multiple comparisons tests: 2x: 95% CI: −0,6449 to −0,1507, p = .001; 3x: 95% CI: −0,8102 to −0,3161, p < .001. Tukey’s multiple comparisons tests: dex: 1x vs. 2x: 95% CI: 0,03337 to 0,5127, p = 0.024)*. *(p ≤ 0.05 (*), p ≤ 0.01 (**), p ≤ 0.001 (***))*. Abbreviations in alphabetical order: AMD1 = adenosylmethionine decarboxylase 1, ASDS = acute social defeat stress, Ctrl = control, dex = dexamethasone, FC = fold change, GAPDH = glyceraldehyde-3-phosphate dehydrogenase, p62 = SQSTM1 = sequestosome 1, put = putrescine, spd = spermidine, veh = vehicle.

A hallmark of stress is the activation of the glucocorticoid receptor (GR) as part of the hypothalamic-pituitary-adrenal axis^12^. To test whether this fundamental stress mechanism could be responsible for the stress-induced changes in polyamine metabolism and consequently in autophagy, we injected mice with the GR antagonist mifepristone (best known as RU486) (mouse #2, Extended Data Fig. 3A),^13^). GR blockade induced alterations of put and AMD1 opposite in direction to those caused by acute stress (Fig. 2D, E) in the brains, livers, and plasma of mice (Extended Data Fig. 3B-F). This indicates that the stress effects on autophagy and polyamine metabolism can be at least partially explained by glucocorticoid signaling. To corroborate these findings, we further treated animals with the selective GR substrate and activator, dexamethasone (dex), using repetitive application to examine potential dose effects (mouse #3, Fig. 2G). In the brain tissue of these animals, we initially assessed the activation of the GR by analyzing the well-established GR target FKBP51, which confirmed a dose-dependent increase in its protein levels (Extended Data Fig. 4A). Repeated administration of this glucocorticoid revealed a dose-dependent increase in put (but not in orn, spd, and spm) (Fig. 2H, Extended Data Fig. 4B), as well as an increase in AMD1 levels with two or three injections compared to one injection (Fig. 2I). These findings are consistent with signs of activated autophagic protein degradation represented by a gradual reduction of p62 in mouse brain tissue, indicating induced autophagy (Fig. 2J).

Overall, polyamine metabolism seems to be highly responsive to acute stressors, particularly affecting put and spd levels, with more pronounced changes observed in peripheral tissues than in the brain. These regulatory effects may be mediated by the GR, as its targeted perturbations show results consistent with those observed in the acute stress response.

### From acute to chronic stress: Altered polyamine dynamics in mouse models of depression and in disease

To address the intriguing question of how the physiological response to acute stressful challenges can turn into a maladaptive pathway that may border on psychopathology and stress-related mental dysfunction, we expanded our studies from acute stress models to a well-established chronic social stress model (CSDS) of depression in mice. We utilized CSDS to further elucidate the role of spd-associated autophagy in stress-associated depressive phenotypes. In our CSDS model, mice underwent daily defeat sessions by a dominant aggressor for 10 days (mouse #4, Fig. 3A). Following this repeated “bullying-like” social stress exposure, mice developed a depression-like phenotype^14^. Tissue samples and plasma were collected one day after the last defeat to investigate the persistent effects of chronic stress exposure (Fig. 3A). Autophagy measurements in the brain, including p62 (Extended Data Fig. 5A), and proteomic analysis (Extended Data Fig. 5B, C, Supplementary Table 4), showed no lasting changes. In contrast, chronic stress led to elevated put levels, as seen previously with acute stressors, but surprisingly also resulted in a notable increase in spd levels in the brain (Fig. 3B, Extended Data Fig. 5D), in line with changes observed in plasma and liver levels (Extended Data Fig. 5E, F). The increase in spd did not lead to enhanced hypusination, possibly explaining the unaltered surrogates of autophagy-mediated protein degradation (Extended Data Fig. 5G, H). At the enzymatic level, AMD1 levels were decreased (Fig. 3C), while ODC1 and SAT1 levels remained constant (Extended Data Fig. 5I, J). It can be speculated that this reduction in AMD1 levels is a response to the chronically elevated spd levels (Fig. 3B). It can be speculated that the reduced AMD1 levels (Fig. 3C) are a response to chronically elevated spd levels (Fig. 3B), while ODC1 and SAT1 levels remain constant (Extended Data Fig. 5I, J).

**Fig. 3.**
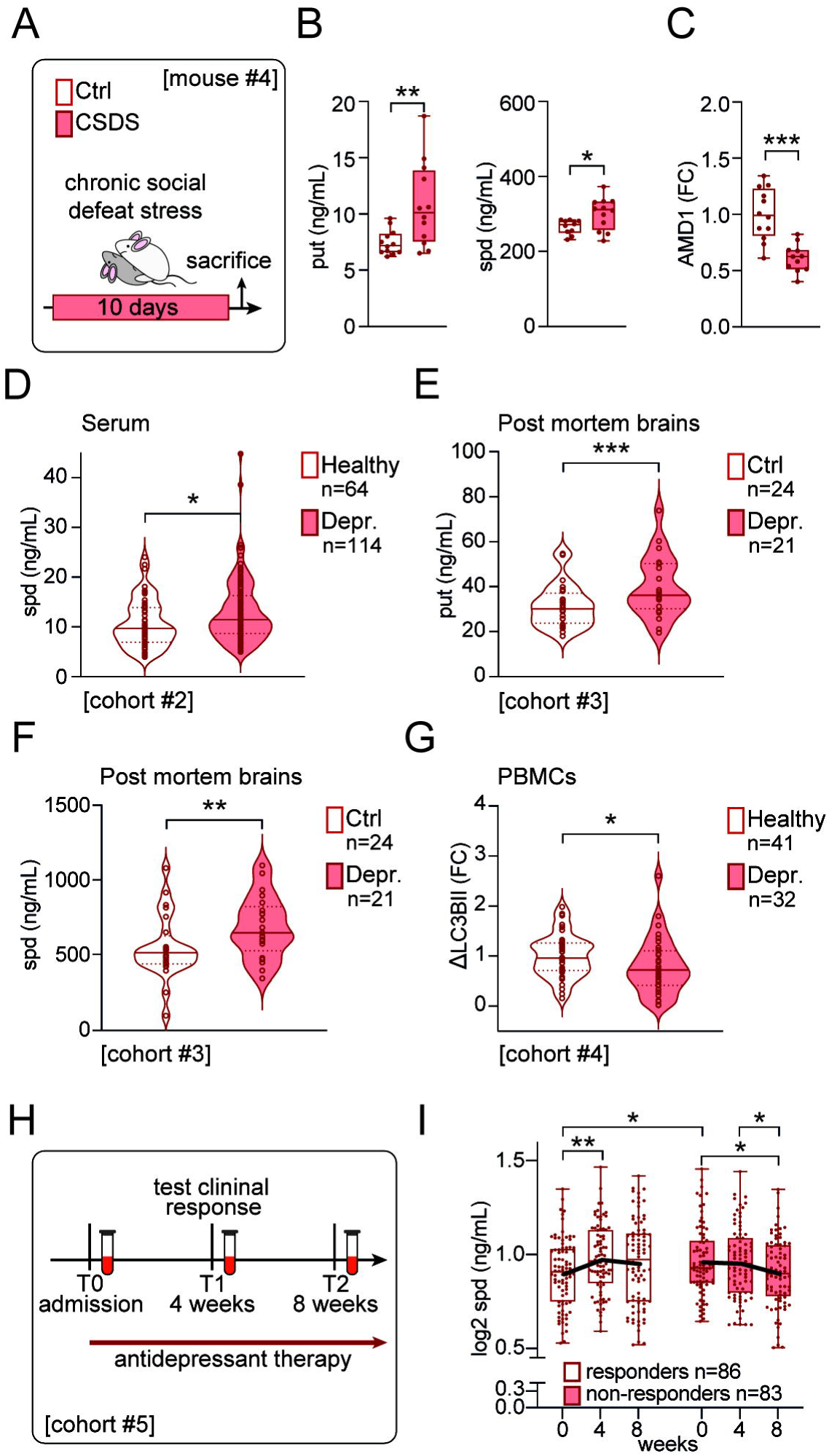
Polyamine levels are altered, and autophagy is reduced in the disease-state. **(A)** Schematic overview of the CSDS paradigm in mice *(mouse #4: n = 12 CSDS, n = 12 Ctrl)*. **(B)** Targeted analysis of put and spd levels in brain lysates from mice in **(A)** (put: n = 12 CSDS, n = 12 Ctrl; unpaired two-tailed t-test: p = 0.0074. spd: n = 12 CSDS, n = 10 Ctrl, unpaired two-tailed t-test: p = 0.0234). **(C)** Relative AMD1 levels in brain lysates from mice in **(A)** determined by immunoblotting *(n = 11 CSDS, n = 12 Ctrl; unpaired two-tailed t-test: p = <0.001)*. **(D)** Targeted analysis of spd levels in serum samples from depressed patients and healthy participants in cohort #2 *(n = 114 Depr., n = 64 Healthy, p value adjusted for significant effects of sex and illegal drug use)*. **(E, F)** Targeted analysis of put and spd levels in postmortem hippocampal brain samples from subjects with depression and control brain donors in cohort #3 *(n = 21 Depr., n = 24 Ctrl,* p value for put *adjusted for significant effect of duration of Depr., p value for spd adjusted* for *significant effect of antidepressant exposure)*. **(G)** Relative ΔLC3B-II levels in PBMCs from depressed patients and healthy participants at baseline, prior to the start of the intervention in cohort #4 *(n = 32 Depr., n = 41 Healthy; Mann-Whitney test: p = 0.0336)*. **(H)** Schematic overview of the study design for cohort #5, involving depressed patients, including the time points for blood sample collection and the duration of antidepressant treatment. **(I)** Targeted analysis of spd in depressed patients from cohort #5, with measurements taken at baseline (T0/BL), after 4 weeks (W4), and after 8 weeks (W8) of antidepressant treatment. The cohort is divided into responders (n = 86) and non-responders (n = 83) according to changes in HAM-D 17 from BL to W4, with the results highlighting the comparison of spd levels between these groups. ANCOVA models reflecting a significant effect of BMI for BL vs. W4, and significant effects of BMI and sex for BL vs. W8 as well as W4 vs. W8 in responders. ANCOVA models for non-responders reflect significant effects of BMI and sex for BL vs. W4. p ≤ 0.05 (*), p ≤ 0.01 (**), p ≤ 0.001 (***) **Abbreviations in alphabetical order:** AMD1 = adenosylmethionine decarboxylase 1, BL = baseline, Ctrl = control, CSDS = chronic social defeat stress, Depr. = depressed, FC = fold change, HAM-D 17 = 17-item Hamilton Depression Rating Scale, LC3B-II = microtubule-associated protein 1 light chain 3 beta-II, PBMCs = peripheral blood mononuclear cells, put = putrescine, spd = spermidine.

To translate our findings from the mouse model to humans, we first assessed polyamine levels in the serum of depressed patients (Supplementary Table 5, cohort #2), revealing an increase in spd, but no changes in orn, put, and spm levels compared to non-depressed controls (Fig. 3D, Extended Data Fig. 6A). When comparing these results with polyamine measurements in postmortem hippocampal brain tissue from depressed patients and controls (Supplementary Table 6, cohort #3), we also observed increased levels of spd, put, and spm, while orn levels remained unaffected (Fig. 3E, F, Extended Data Fig. 6B). Especially the effect of increased spd levels appears to be very robust, even in depressed patients with comorbid substance use disorder, showing elevated spd and put levels (Extended Data Fig. 6C). Thus, the transient activation of autophagy via hypusination seems to be lost during chronic stress. As a consequence, the organism tries to compensate via spd production in a maladaptive fashion.

In line, there is also an increase in spd and put when comparing individuals with substance use disorder to controls, suggesting a role for polyamines across different categories of mental disorders (Extended Data Fig. 6D). To assess the disease-related effects of depression on autophagic flux, we conducted the previously described *ex vivo* autophagy assay on PBMCs from both drug-naive depressed patients and healthy controls (Supplementary Table 7, cohort #4). Accordingly, we observed reduced autophagic flux in depressed patients compared to healthy controls (Fig. 3G, Extended Data Fig. 6E). Interestingly, within this cohort, autophagic flux was positively correlated with resilience scores and well-being scores (Warwick-Edinburgh Mental Wellbeing Scale (WEMWBS)) (Extended Data Fig. 6F, G).

In a prior study, we and others demonstrated a correlation between the clinical response of psychiatric patients to antidepressants and the induction of autophagy, measured *ex vivo* in circulating leukocytes^15^. To investigate whether polyamines play a crucial role in the effectiveness of antidepressants, we measured changes in polyamine levels in the plasma of 169 depressed patients undergoing psychopharmacotherapy (Supplementary Table 8, cohort #5). Plasma was collected before the start of treatment (T0), as well as after 4 (T1) and 8 weeks (T2) of antidepressant treatment (Fig. 3H). Clinical response was assessed after 4 weeks using various established questionnaires and evaluations by the treating physician. In the group of treatment responders, we observed an increase in spd levels, whereas a clear reduction in its levels was noted in the non-responder group over the course of the treatment (Fig. 3I). Orn and put did not show a comparable progression (Extended Data Fig. 6H). Interestingly, non-responders exhibited higher spd levels compared to responders before treatment, suggesting an exhausted feedback mechanism with limited systemic capacity to further elevate spd levels in response to depression, potentially explaining their lack of treatment response (Fig. 3I).

Overall, polyamine metabolism is highly responsive to chronic stressors, showing notably increased spd levels, in contrast to acute stress, which causes an attenuation in spd levels. In depressed, drug-naive patients, autophagic flux was found to be reduced compared to healthy controls. Additionally, our findings suggest that a polyamine-centered metabolic response, characterized by increased circulating spd levels, is essential for the antidepressant effects of standard therapies.

### Spermidine administration alleviates depression in mice and humans

We next investigated whether the successful treatment response to a conventional antidepressant, which was marked by increasing spd levels, could be replicated, through exogenous intraperitoneal (i.p.) SPD supplementation in a mouse model of depression.

To test this hypothesis, mice were given daily i.p. injections of SPD for 26 days, with 3 weeks of CSDS exposure beginning on day 8 (mouse #5, Fig. 4A). Behavioral tests (open field test, social avoidance test, and home cage social interaction test) were conducted in the third week of CSDS, prior to daily defeats. On day 27, all mice were sacrificed in the morning, 24 hours after the last defeat and SPD injections, and sample collections for metabolite and protein analyses were conducted (Fig. 4A). Impaired social behavior, a key feature of MDD and the CSDS paradigm, was observed as increased social avoidance in vehicle-treated CSDS mice but was alleviated by i.p. SPD treatment in the social avoidance test (Fig. 4B) and confirmed in the home cage social interaction test (Fig. 4C). Additionally, while chronic stress led to an overall increase in adrenal gland weight, i.p. SPD treatment resulted in lower adrenal gland weight, regardless of stress condition (Extended Data Fig. 7B).

**Fig. 4.**
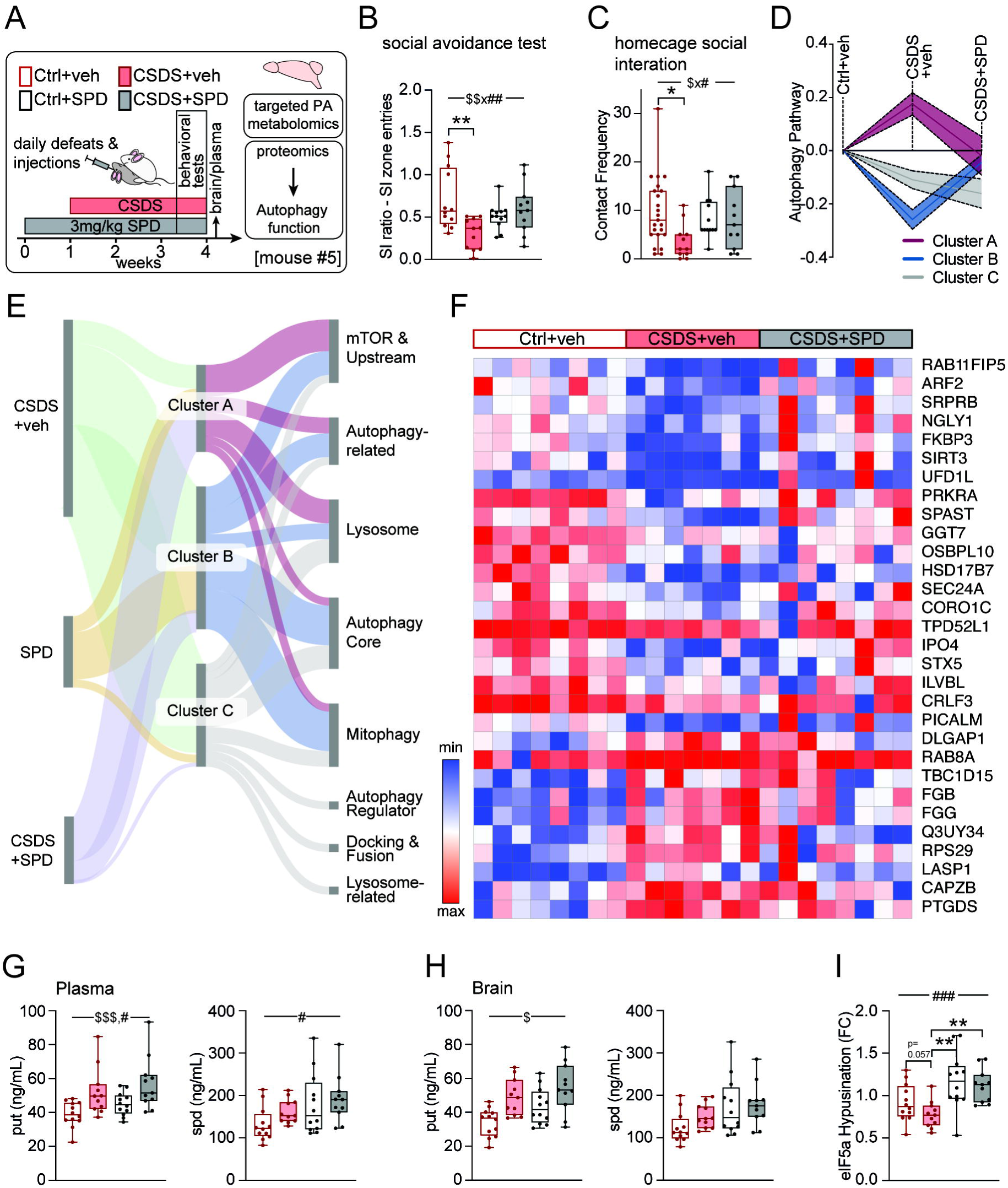
SPD treatment alleviates depressive-like symptoms and counteracts stress-induced alterations in autophagy-related protein levels in mice. **(A)** Schematic overview of the study design involving four groups of mice: two groups subjected to CSDS with either SPD or veh injections (intraperitoneal, i.p.), and two control groups receiving either SPD or veh i.p. along with an overview of subsequent tests and analyses (mouse #5, *n = 12 Ctrl+veh, n = 12 Ctrl+SPD, n = 11 CSDS+SPD, n = 11 CSDS+veh)*. **(B)** Social interaction ratio was measured in a social avoidance test from mice in **(A)** (SI ratio - SI zone entries: n = 12 Ctrl+veh, n = 12 Ctrl+SPD, n = 11 CSDS+SPD, n = 11 CSDS+veh; two-way ANOVA: CSDS x SPD interaction: F_(1, 42)_ = 10.19, p = 0.0027. Tukey’s multiple comparisons test: Ctrl+veh vs. CSDS+veh: 95% CI: 0.1139 to 0.7041, p = 0.0033). **(C)** Contact frequency was assessed in a home cage social interaction test from mice in **(A)** (contact frequency: n = 12 Ctrl+veh, n = 12 Ctrl+SPD, n = 11 CSDS+SPD, n = 11 CSDS+veh; two-way ANOVA, CSDS x SPD interaction: F_(1, 42)_ = 4.834, p = 0.0305. Tukey’s multiple comparisons test: Ctrl+veh vs. CSDS+veh: 95% CI: 1.099 to 13.79, p = 0.0158). **(D)** Line plot displaying the expression levels of clusters of autophagy-related proteins in mice from **(A)** *(n = 8 Ctrl+veh, n = 8 Ctrl+SPD, n = 8 CSDS+SPD, n = 7 CSDS+veh)*. **(E)** Sankey plot illustrating clusters of autophagy-associated proteins altered by CSDS, SPD, or the interaction between these factors, and their categorization *(n = 8 Ctrl+veh, n = 8 Ctrl+SPD, n = 8 CSDS+SPD, n = 7 CSDS+veh)*. **(F)** Heatmap illustrating the expression patterns of the top 30 altered proteins due to interventions from **(A)** *(n = 8 Ctrl+veh, n = 8 Ctrl+SPD, n = 8 CSDS+SPD, n = 7 CSDS+veh)*. **(G)** Targeted analysis of put and spd in plasma samples from mice in **(A)** (Plasma: n = 12 Ctrl+veh, n = 12 Ctrl+SPD, n = 11 CSDS+SPD, n = 11 CSDS+veh; two-way ANOVA: put: main CSDS effect: F_(1, 42)_ = 15.39, p = 0.0003, main SPD effect: F_(1, 42)_ = 5.316, p = 0.0261. spd: main SPD effect: F_(1, 42)_ = 7.111, p = 0.0108). **(H)** Targeted analysis of put and spd in brain lysates from mice in **(A)** (*Brain: n = 12 Ctrl+veh, n = 12 Ctrl+SPD, n = 11 CSDS+SPD, n = 10 CSDS+veh; put: two-way ANOVA: main CSDS effect: F_(1, 41)_ = 5.955, p = 0.0191)*. **(I)** Relative levels of hypusinated eIf5A in brain samples from mice in **(A)** determined by immunoblotting *(n = 12 Ctrl+veh, n = 12 Ctrl+SPD, n = 11 CSDS+SPD, n = 11 CSDS+veh; two-way ANOVA: main SPD^-^effect: F_(1, 42)_ = 17,72, p <.001. Tukey’s multiple comparisons test: Ctrl+SPD vs CSDS+veh: 95% CI: 0,1317 to 0,6640, p = 0.001. CSDS+veh vs. CSDS+SPD: - 0,6091 to −0,06541, p = 0.010. Ctrl+veh vs. Ctrl+SPD: 95% CI:* −0,5153 to 0,005293*, p = 0.057).* p ≤ 0.05 (*), p ≤ 0.01 (**), p ≤ 0.001 (***) **Abbreviations in alphabetical order:** Ctrl = control, CSDS = chronic social defeat stress, eIF5A = eukaryotic translation initiation factor 5A, FC = fold change, PA = polyamine, SPD = spermidine i.p. injected, spd = spermidine (metabolite), $ = CSDS effect, # = SPD effect.

Not all features of depression were alleviated by spermidine, as i.p. administration of SPD did not affect anxiety-related behavior in the open field test (Extended Data Fig. 7A).

To comprehensively assess the impact of various stimuli and their combination on the autophagy machinery, we conducted a proteomic analysis of whole-brain tissue from mice (Extended Data Fig. 7C, Supplementary Table 9). We observed changes in the expression of proteins due to CSDS, treatment, or the combination of both. Consequently, these proteins were classified based on their involvement as components of the autophagy pathway, according to expert-curated lists^16^. These autophagy-related proteins were then ranked into three clusters based on their expression patterns (Fig. 4D). Clusters A and B, representing 69% of the altered autophagy-related proteins (29 out of 42), showed elevated or decreased expression, respectively, in the vehicle-treated CSDS group compared to controls. Notably, expression levels of these clusters in the i.p. SPD-treated CSDS group were similar to those of the control group, indicating a potential effect of SPD in counteracting stress-induced alterations in autophagy-related protein levels. Cluster A proteins are primarily involved in the mTOR signaling pathway and upstream regulatory pathways (Pik3ca and Atp6v1g1), and lysosomal functions (Ctsb and Vps18), while Cluster B proteins are more involved in the core autophagy machinery (LC3B and Picalm) and mitophagy (Fis1 and Htra2) (Fig. 4E, F).

Next, we determined the levels of polyamines in the brain, plasma, and liver. Consistent with our previous findings, we observed strong effects of CSDS on put and spd levels in the periphery, e.g. in plasma and liver (Fig. 4G, Extended Data Fig. 7E). SPD treatment had the most pronounced effects on peripheral tissues, particularly the liver and plasma (Fig. 4G, Extended Data Fig. 7E, F). The brain appeared to undergo less pronounced fluctuations in this experimental design (Fig. 4H, Extended Data Fig. 7D). Additionally, in mouse brain tissue, we assessed the level of eIF5A hypusination and observed that i.p. treatment with SPD led to a significant increase in hypusination in CSDS animals (Fig. 4I).

In summary, this experiment clearly demonstrates that i.p. SPD supplementation is able to counterbalance and restore chronic social stress-induced behavioral abnormalities, comparable to the effects of antidepressants. We hypothesize that the antidepressant-like effects are mediated through SPD-induced restoration of autophagy levels under chronic stress conditions.

Based on the preclinical results suggesting an antidepressant-like effect of SPD, we conducted a double-blind, placebo-controlled clinical study (Fig. 5A, Extended Data Fig. 8A, Supplementary Table 7, cohort #4). 44 healthy volunteers and 35 drug-naive mildly to moderately depressed patients were administered SPD daily (6 mg per day through a spermidine-rich wheat germ extract) or a placebo for a total of three weeks (Fig. 5A). The participants were assessed at baseline (T0) and at weekly intervals, including a follow-up session (T4) one week after the end of treatment.

**Fig. 5.**
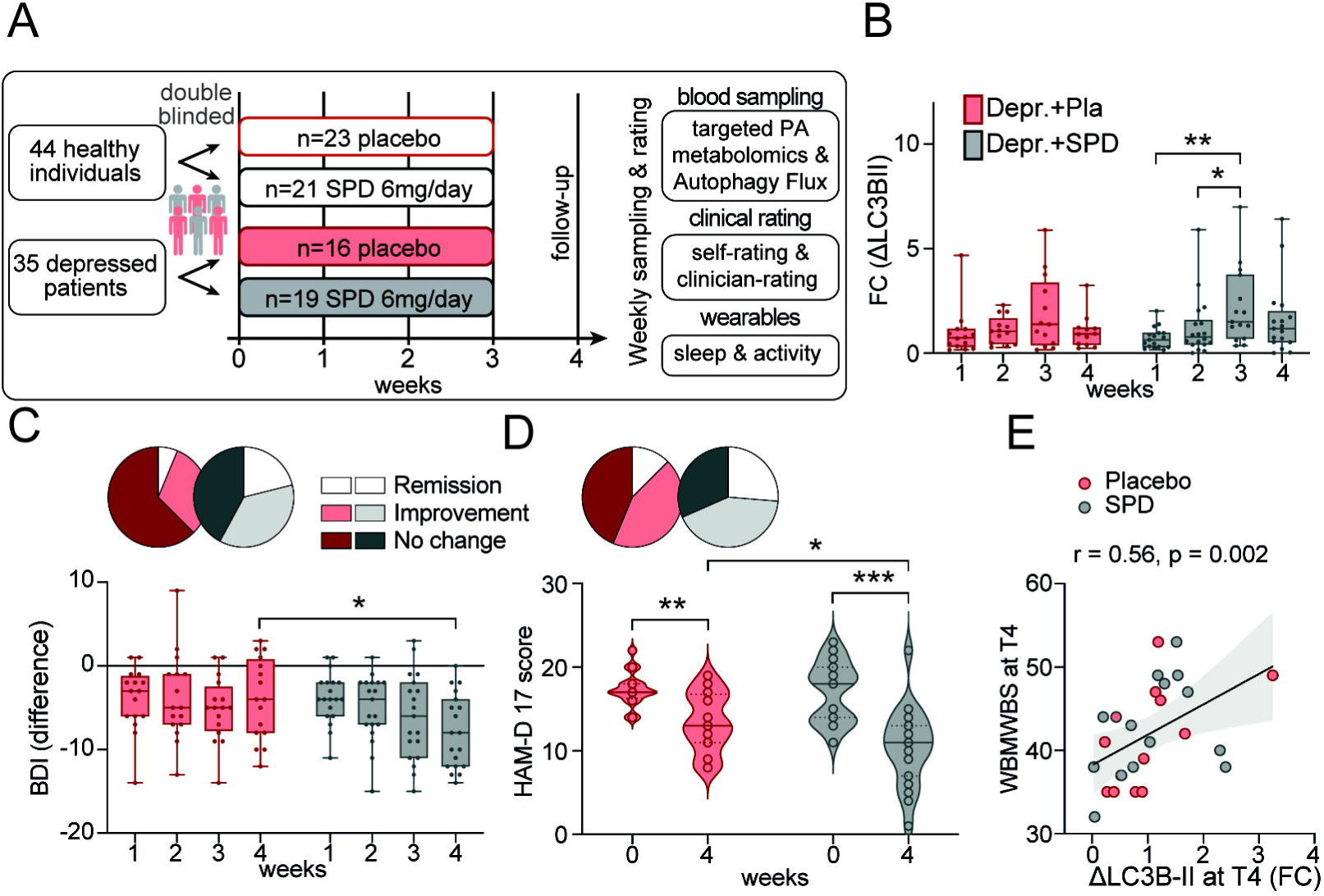
SPD supplementation alleviates depressive symptoms and induces autophagy in humans. **(A)** Schematic overview of the double-blind, placebo-controlled, randomized study design for cohort #4, including SPD or placebo supplementation, clinical assessments, wearables, and blood sample collection, followed by subsequent blood analyses (*n = 35 Depr., n = 44 healthy individuals*). **(B)** Relative ΔLC3B-II levels in PBMCs from depressed patients in cohort #4, assessing the effects of SPD^+^ and Pla throughout the study *(n = 18 SPD, n = 14 Pla; Sidak’s multiple comparisons test: T1 vs. T3: 95% CI: −0.7449 to −0.1556, hedges’ g = 1.1352, p = 0.0019; T2 vs. T3: 95% CI: −0.5407 to −0.02758, p = 0.0249, hedges’ g = 0.5918)*. **(C)** Analysis of the BDI presented as difference scores in depressed patients receiving SPD or placebo supplementation from cohort #4 throughout the study. Pie charts illustrate the rates of remission, improvement, and no change in symptom severity: Remission is defined as a score of less than 10, improvement as a decrease of 8 points or more, and no change indicates stability in symptom severity *(Remission: SPD: 4 out of 19, Pla: 1 out of 16; improvement: SPD: 8 out of 19, Pla: 2 out of 16; no change: SPD: 1 out of 19, Pla: 5 out of 16. Week 4: SPD vs. Pla: Sidak’s multiple comparisons test: 95% CI: 0.2459 to 7.669, p = 0.0316, hedges’ g = 0.8559)*. **(D)** Analysis of the HAM-D 17 in depressed patients receiving SPD or placebo supplementation from cohort #4 at baseline and follow-up appointments. Pie charts illustrate the rates of remission, improvement, and no change in symptom severity: Remission is defined as a score of 8 or less, improvement is defined as a decrease of 8 points or more, and no change indicates stability in symptom severity (*Remission: SPD: 5 out of 19, Pla: 2 out of 16; improvement: SPD: 7 out of 19, Pla: 4 out of 16; no change: SPD: 3 out of 19, Pla: 4 out of 16. Sidak’s multiple comparisons test: Pla: Week 0 vs. Week 4: 95% CI: 1.276 to 6.599, p = 0.0030, hedges’ g = 1.3703; SPD: Week 0 vs. Week 4: 95% CI: 4.189 to 9.074, p = < 0.001, hedges’ g = 1.5795; Pla vs. SPD Week 4: 95% CI: 0.1249 to 5.803, p = 0.0391, hedges’ g = 0.7029)*. **(E)** Correlation analysis between relative ΔLC3B-II levels and WEMWBS in depressed patients from cohort #4 at follow-up appointment *(n = 15 SPD, n = 12 Pla, Spearman’s rank correlation: r = 0.56, p = 0.002).* p ≤ 0.05 (*), p ≤ 0.01 (**), p ≤ 0.001 (***) **Abbreviations in alphabetical order:** BDI = Beck Depression Inventory, FC = fold change, HAM-D 17 = 17-item Hamilton Depression Rating Scale, LC3B-II = microtubule-associated protein 1 light chain 3 beta-II, Pla = placebo, SPD = spermidine (supplement), WEMWBS = Warwick-Edinburgh Mental Wellbeing Scale.

Initially, we assessed the autophagic flux in participants’ blood and observed that SPD supplementation led to an increase in week 3 of the SPD treatment (Fig. 5B, Extended Data Fig. 8B). However, orn or polyamine plasma levels did not change upon SPD supplementation (Extended Data Fig. 8C), confirming previous reports that blood polyamine levels are not indicative of SPD treatments in humans^17^.

Of note, depressed patients treated with SPD demonstrated clinical improvement, as assessed by changes from baseline in Beck Depression Inventory (BDI) and Hamilton Depression Rating Scale (HAM-D 17) (Fig. 5C, D). Interestingly, autophagic flux showed a positive correlation with the WEMWBS (Fig. 5E). Moreover, the ratio of responders to non-responders, as assessed by BDI and HAM-D 17 scores, clearly indicates that SPD-treated patients benefited more from the intervention compared to the placebo group (Extended Data Fig. 8D).

We concluded that SPD supplementation alleviates depressive symptoms as shown in two different depression scales and increases autophagic flux *in vivo* in PBMCs of depressed patients.

## Discussion

Developing novel therapeutic options for depression is crucial, as current lack efficiency and are prone to side-effects. Despite compelling evidence that SRPDs are systemic, most therapies remain brain-focused. Our study highlights the significant involvement of polyamine metabolism and its interaction with autophagy in acute stress and SRPDs, both in mice and humans. We identified that acute and chronic stress modulate polyamine levels throughout the body, having found metabolic changes in plasma and postmortem brain tissues of humans, as well as in the plasma, brain and peripheral tissues of mice. Further, we uncovered a mechanistic involvement of glucocorticoids and GR signaling in this shift in polyamine metabolism, in a dose-dependent manner.

We linked autophagy dysregulation, including disruptions in the polyamine cascade and reduced autophagic flux to SRPDs such as depression, which aligns with our findings from a mouse model of depression. Furthermore, an endogenous increase in spd plasma levels, likely resulting from activated biosynthesis during antidepressant pharmacotherapy, could have the potential to predict clinical treatment response. For the first time, we show that the i.p. administration of SPD improves depressive-like behavior in mice and restores autophagy. Moreover, in a double-blind, placebo-controlled trial involving drug-naive depressed patients, SPD supplementation led to improvements in depressive symptoms and an increase in autophagic flux.

Given these findings, along with the observation that non-responders had higher spd levels compared to responders before treatment, further studies should investigate whether SPD supplementation could restore the effectiveness of antidepressant therapies. This would support the hypothesis that non-responders may have an exhausted feedback mechanism with a limited capacity to further increase spd production in response to depression.

### Acute stress affects autophagy and related metabolic pathways

Using acute stress models in humans and mice, we aimed to elucidate the impact of a single stressful challenge on autophagy and its regulatory pathways in cellular metabolism across the brain and body. On the mechanistic level, a particular focus has been placed on glucocorticoid signaling. Previous studies suggested that acute cellular stress triggers autophagy, most likely via glucocorticoid signaling^18^, assisting cells in eliminating damaged components, thereby enhancing survival and ensuring optimal function^19^. We found an increased autophagic flux using a tailored *ex vivo* assay, before and after a bungee jump, in parallel with rising cortisol levels. This anticipatory stress before the bungee jump suggests that psychological stress can elevate autophagic flux^8^. Additionally, a similar increase in autophagy was observed in acutely stressed mice using a defined, single social stressor. Glucocorticoid administration or antagonization of the GR in mice demonstrated that stress-induced changes in autophagy and polyamine levels are in crosstalk with GR signaling. This aligns with studies showing that dex treatment leads to an increase in put levels^20^ and autophagy signaling^15^. Data from published mouse studies support our findings on the connection between polyamine levels and acute stress, showing consistent regulation of polyamines as part of the stress response. Acute stress in animal models increased put levels in the brain and periphery^21^, while spd and spm levels remained largely unchanged^22^. The bungee jump stress model in humans mainly showed changes in precursor metabolites such as arg and cit of the polyamine pathway and depletion of spd, indicating that different stressors might affect different stages of the cascade based on its intensity and duration. The regulation of the polyamine cascade appears to be significantly influenced by consistently elevated levels of AMD1 due to acute stress. In addition to glucocorticoids, as previously discussed, the main polyamine antizymes and synthetases exhibit only a minor change in response to stress.

Taken together, our bungee jump experiment demonstrated that a single, powerful stressor - combining both psychological and physical elements and involving various physiological mechanisms - alters polyamine levels and autophagy, with GR signaling playing a role. Intriguingly, we were able to replicate these findings in a mouse model using a social stressor, suggesting that this mechanism is fundamental across different types of stressors.

### The polyamine hypothesis of depression

Aberrant polyamine signaling has previously been linked to the pathogenesis of depression and other SRPDs, although this connection has remained somewhat uncertain. For instance, it has been shown that the depletion of the spd precursor put by the ODC1 antagonist DFMO exhibited depressogenic effects in rats^23^. However, our current data from both mouse models of depression and independent human cohorts, now provide strong, converging evidence that disturbances in polyamine metabolism are indeed involved in the development of depression.

In alignment with the shift from depleted spd during acute stress to an increase during chronic stress, we observed a pronounced downregulation of AMD1 specifically during chronic stress in the brain. This reduction in AMD1 may result in elevated s-adenosylmethionine (SAM) levels and impaired conversion to 5’-deoxy-5’-methylthioadenosin (dcSAM)^24^, consistent with effects seen in AMD1 knockdown models or with inhibitors^25^. SAM is crucial for DNA methylation and is commonly elevated in depression, while dcSAM inhibits DNA methyltransferases. Therefore, reduced AMD1 levels and the resulting changes in SAM:dcSAM ratios could engender epigenetic changes associated with depression. Since SAM is crucial for DNA methylation and is commonly elevated in depression, while dcSAM inhibits DNA methyltransferases, reduced AMD1 levels could promote epigenetic changes associated with depression. Thus, AMD1 may function as a molecular switch linking the autophagy-promoting polyamine pathway to the epigenetic mechanisms underlying depressive disorders^26,27^.

Our analyses of human cohorts with depression confirmed distinct polyamine profiles and reduced autophagic flux, along with significant alterations in plasma spd levels, indicating a specific involvement of polyamines and autophagy in the pathophysiology of depression. These findings were validated in postmortem hippocampal brain samples, consistent with previous research on depressed patients^28^ and suicide completers^29^. These disturbances in the polyamine-autophagy axis affect both peripheral and central organs, supporting the view of depression as a systems biology disease. Moreover, we showed that increased spd levels during depression treatment are essential for achieving clinical success and may also serve as a potential biomarker of treatment response. Understanding the mechanistic insights behind why some people respond to pharmacotherapy, while others are resistant could guide treatment decisions and open avenues for the development of innovative therapeutic strategies.

Our clinical study suggests that SPD, which elevates autophagic flux, is a promising treatment option for depression, showing effect sizes larger than those of approved antidepressant therapies^30^. Specifically, BDI scores reveal a large effect size (Hedges’ g = 0.85) for SPD treatment compared to placebo at Week 4, which is substantially higher than the typical effect size of 0.30 to 0.40 reported in meta-analyses of antidepressants for depression^31^. Similarly, the HAM-D 17 scores reveal a moderate effect size when comparing the placebo group with the SPD treatment group at Week 4 (Hedges’ g = 0.70). Within-group comparisons of HAM-D 17 scores further support this trend, with large effect sizes observed (Hedges’ g = 1.37 for placebo and 1.58 for SPD). SPD intake is commonly associated with life extension and neuroprotection and has been linked to a reduced prevalence of depressive symptoms in observational studies^32^. Participants in our study had mild to moderate depressive symptoms, which may limit the generalizability of our findings to more severe cases. Nevertheless, given that antidepressant medications are often more effective for severe depressive episodes^33^, SPD supplementation might also benefit this group, potentially in synergy with existing antidepressant therapies. This possibility should be further investigated in future clinical trials to assess its validity and therapeutic potential.

Overall, our study underscores the pivotal role of polyamine metabolism, in conjunction with autophagy, as a crucial mechanistic link to stress. Polyamine-mediated autophagy, potentially mediated by glucocorticoid signaling pathways, plays a significant role in the acute stress response, affecting energy balance and protein homeostasis. In depression, this axis appears to be dysregulated, leading to reduced autophagic flux and a substantial dysregulation of polyamine signaling, which might promote DNA methylation through SAM accumulation. This can result in a rigid chromatin structure and contribute to disease progression. Additionally, the inducibility of spd is critical for the responsiveness to antidepressants, as demonstrated by one of our clinical studies. Reactivating autophagy is crucial for recovery from SRPDs. This is particularly relevant, as it has been shown that repeatedly stressed mice adopt a starvation-like metabolism^34^, with starvation being a primary stimulator of autophagy. Notably, we recently demonstrated that starvation requires the polyamine cascade to fully initiate autophagy and to exert its cell-protective homeostatic effects^35^.

In summary, our study reveals that the polyamine-autophagy axis is a crucial mechanism in both acute and chronic stress response and represents a novel therapeutic target for SRPDs and depression.

## Data availability

The mass spectrometry proteomics data have been deposited to the ProteomeXchange Consortium via the PRIDE^36^ partner repository with the dataset identifiers PXD054338 (Mouse brains after CSD treated with SPD or vehicle), PXD053802 (Mouse brains after CSD), PXD053796 (Mouse brains after ASD). All other data supporting the findings of this study are available from the corresponding author on reasonable request.

## Code availability

For this manuscript no custom algorithm or software was used.

## Acknowledgements

We thank all lab members for suggestions and comments on the experiments and manuscript. We are grateful for the technical assistance provided by M. Hausl (Joanneum Research HEALTH, Graz, Austria).

## Author contributions

Conceptualisation: SM, CN, TEbert, FM, NCG

Methodology: SM, CN, TEbert, TB, SD, AZ, BG, DEH, AMüller, LO, AMeinitzer, HP, MK, MBM, TEisenberg, JH, NCG

Software: TEbert, SD

Validation: BS-W, BK, KJR, GK, TEisenberg, JK, FM, NCG

Formal analysis: SM, CN, YM, TEbert, TB, SD, SJH, AZ, ELN, AMeinitzer, HP, TEisenberg, JH, NCG

Investigation: SM, CN, YM, TEbert, TB, SD, SJH, AZ, JE, VK, LAH, MLennarz, ELN, CS, DEH, EJ, CK, AMeinitzer, MLaakmann, RA, MB, KL, BS-W, NO, MVS, HP, MK, MBM, TEisenberg, JH, NCG

Resources: BG, AMeinitzer, AP, HR, BS-W, BK, KJR, HP, MBM, GK, TEisenberg, JH, NCG Data Curation: SM, CN, YM, TB, SD, SJH, AZ, BB, ELN, TEisenberg, JH, NCG

Writing - Original Draft: SM, YM, NCG

Writing - Review & Editing: CN, TEbert, TB, SJH, NO, MVS, GK, TEisenberg, FM Visualization: YM, TB, NCG

Supervision: SM, CN, BK, GK, JH, TEisenberg, FM, NCG Project administration: JH, FM, GK, FM, NCG

Funding acquisition: SM, TEbert, BB, NCG

## References

1. Akil H, Nestler EJ. The neurobiology of stress: Vulnerability, resilience, and major depression. Proc Natl Acad Sci U S A. 2023;120(49):e2312662120. doi:10.1073/pnas.2312662120

2. LeMoult J. From Stress to Depression: Bringing Together Cognitive and Biological Science. Curr Dir Psychol Sci. 2020;29(6):592–598. doi:10.1177/0963721420964039

3. Marx W, Penninx BWJH, Solmi M, et al. Major depressive disorder. Nat Rev Dis Primer. 2023;9(1):44. doi:10.1038/s41572-023-00454-1

4. Correll CU, Solmi M, Cortese S, et al. The future of psychopharmacology: a critical appraisal of ongoing phase 2/3 trials, and of some current trends aiming to de-risk trial programmes of novel agents. World Psychiatry Off J World Psychiatr Assoc WPA. 2023;22(1):48–74. doi:10.1002/wps.21056

5. Wittenborn AK, Rahmandad H, Rick J, Hosseinichimeh N. Depression as a systemic syndrome: mapping the feedback loops of major depressive disorder. Psychol Med. 2016;46(3):551–562. doi:10.1017/S0033291715002044

6. Levine B, Kroemer G. Autophagy in the pathogenesis of disease. Cell. 2008;132(1):27–42. doi:10.1016/j.cell.2007.12.018

7. Floriou-Servou A, von Ziegler L, Waag R, Schläppi C, Germain PL, Bohacek J. The Acute Stress Response in the Multiomic Era. Biol Psychiatry. 2021;89(12):1116–1126. doi:10.1016/j.biopsych.2020.12.031

8. Ebert T, et al. Acute stress triggers RNA splicing and protein secretion. submitted.

9. Zahedi K, Barone S, Soleimani M. Polyamines and Their Metabolism: From the Maintenance of Physiological Homeostasis to the Mediation of Disease. Med Sci Basel Switz. 2022;10(3):38. doi:10.3390/medsci10030038

10. Bensalem J, Hattersley KJ, Hein LK, et al. Measurement of autophagic flux in humans: an optimized method for blood samples. Autophagy. Published online December 11, 2020:1–18. doi:10.1080/15548627.2020.1846302

11. Uribe-Mariño A, Gassen NC, Wiesbeck MF, et al. Prefrontal Cortex Corticotropin-Releasing Factor Receptor 1 Conveys Acute Stress-Induced Executive Dysfunction. Biol Psychiatry. 2016;80(10):743–753. doi:10.1016/j.biopsych.2016.03.2106

12. Nicolaides NC, Charmandari E, Chrousos GP, Kino T. Circadian endocrine rhythms: the hypothalamic-pituitary-adrenal axis and its actions. Ann N Y Acad Sci. 2014;1318:71–80. doi:10.1111/nyas.12464

13. Castinetti F, Conte-Devolx B, Brue T. Medical treatment of Cushing’s syndrome: glucocorticoid receptor antagonists and mifepristone. Neuroendocrinology. 2010;92 Suppl 1:125–130. doi:10.1159/000314224

14. Golden SA, Covington HE, Berton O, Russo SJ. A standardized protocol for repeated social defeat stress in mice. Nat Protoc. 2011;6(8):1183–1191. doi:10.1038/nprot.2011.361

15. Gassen NC, Hartmann J, Zschocke J, et al. Association of FKBP51 with priming of autophagy pathways and mediation of antidepressant treatment response: evidence in cells, mice, and humans. PLoS Med. 2014;11(11):e1001755. doi:10.1371/journal.pmed.1001755

16. Bordi M, De Cegli R, Testa B, Nixon RA, Ballabio A, Cecconi F. A gene toolbox for monitoring autophagy transcription. Cell Death Dis. 2021;12(11):1044. doi:10.1038/s41419-021-04121-9

17. Senekowitsch S, Wietkamp E, Grimm M, et al. High-Dose Spermidine Supplementation Does Not Increase Spermidine Levels in Blood Plasma and Saliva of Healthy Adults: A Randomized Placebo-Controlled Pharmacokinetic and Metabolomic Study. Nutrients. 2023;15(8):1852. doi:10.3390/nu15081852

18. Martinelli S, Anderzhanova EA, Wiechmann S, et al. Stress-primed secretory autophagy drives extracellular BDNF maturation. bioRxiv. Published online January 1, 2020:2020.05.13.090514. doi:10.1101/2020.05.13.090514

19. Li W, He P, Huang Y, et al. Selective autophagy of intracellular organelles: recent research advances. Theranostics. 2021;11(1):222–256. doi:10.7150/thno.49860

20. Ientile R, De Luca G, Di Giorgio RM, Macaione S. Glucocorticoid regulation of spermidine acetylation in the rat brain. J Neurochem. 1988;51(3):677–682. doi:10.1111/j.1471-4159.1988.tb01797.x

21. Gilad GM, Gilad VH. Polyamine biosynthesis is required for survival of sympathetic neurons after axonal injury. Brain Res. 1983;273(1):191–194. doi:10.1016/0006-8993(83)91113-7

22. Gilad GM, Gilad VH. Overview of the brain polyamine-stress-response: regulation, development, and modulation by lithium and role in cell survival. Cell Mol Neurobiol. 2003;23(4-5):637–649. doi:10.1023/a:1025036532672

23. Gupta N, Zhang H, Liu P. Behavioral and neurochemical effects of acute putrescine depletion by difluoromethylornithine in rats. Neuroscience. 2009;161(3):691–706. doi:10.1016/j.neuroscience.2009.03.075

24. Soda K. Spermine and gene methylation: a mechanism of lifespan extension induced by polyamine-rich diet. Amino Acids. 2020;52(2):213–224. doi:10.1007/s00726-019-02733-2

25. Rahim AB, Lim HK, Tan CYR, et al. The Polyamine Regulator AMD1 Upregulates Spermine Levels to Drive Epidermal Differentiation. J Invest Dermatol. 2021;141(9):2178–2188.e6. doi:10.1016/j.jid.2021.01.039

26. Nestler EJ. Epigenetic mechanisms of depression. JAMA Psychiatry. 2014;71(4):454–456. doi:10.1001/jamapsychiatry.2013.4291

27. Tsankova N, Renthal W, Kumar A, Nestler EJ. Epigenetic regulation in psychiatric disorders. Nat Rev Neurosci. 2007;8(5):355–367. doi:10.1038/nrn2132

28. Wingo TS, Liu Y, Gerasimov ES, et al. Brain proteome-wide association study implicates novel proteins in depression pathogenesis. Nat Neurosci. 2021;24(6):810–817. doi:10.1038/s41593-021-00832-6

29. Chen GG, Fiori LM, Moquin L, et al. Evidence of altered polyamine concentrations in cerebral cortex of suicide completers. Neuropsychopharmacol Off Publ Am Coll Neuropsychopharmacol. 2010;35(7):1477–1484. doi:10.1038/npp.2010.17

30. Khan A, Fahl Mar K, Faucett J, Khan Schilling S, Brown WA. Has the rising placebo response impacted antidepressant clinical trial outcome? Data from the US Food and Drug Administration 1987-2013. World Psychiatry Off J World Psychiatr Assoc WPA. 2017;16(2):181–192. doi:10.1002/wps.20421

31. Hieronymus F, Lisinski A, Eriksson E, Østergaard SD. Do side effects of antidepressants impact efficacy estimates based on the Hamilton Depression Rating Scale? A pooled patient-level analysis. Transl Psychiatry. 2021;11(1):249. doi:10.1038/s41398-021-01364-0

32. Qi G, Wang J, Chen Y, Wei W, Sun C. Association between dietary spermidine intake and depressive symptoms among US adults: National Health and Nutrition Examination Survey (NHANES) 2005-2014. J Affect Disord. 2024;359:125–132. doi:10.1016/j.jad.2024.05.041

33. Fournier JC, DeRubeis RJ, Hollon SD, et al. Antidepressant drug effects and depression severity: a patient-level meta-analysis. JAMA. 2010;303(1):47–53. doi:10.1001/jama.2009.1943

34. Kucukdereli H, Amsalem O, Pottala T, et al. Repeated stress triggers seeking of a starvation-like state in anxiety-prone female mice. Neuron. 2024;112(13):2130–2141.e7. doi:10.1016/j.neuron.2024.03.027

35. Hofer SJ, Daskalaki I, Bergmann M, et al. Spermidine is essential for fasting-mediated autophagy and longevity. Nat Cell Biol. 2024;26(9):1571–1584. doi:10.1038/s41556-024-01468-x

36. Perez-Riverol Y, Bai J, Bandla C, et al. The PRIDE database resources in 2022: a hub for mass spectrometry-based proteomics evidences. Nucleic Acids Res. 2022;50(D1):D543–D552. doi:10.1093/nar/gkab1038

