## Supplementary Information for "Spermidine alleviates depression via control of the stress response"

**Extended Data**

Extended Data Figures 1-8 and legends

**Supplementary Information**

Supplementary Tables 1-9 and legends

Source Data

Material and Methods

### Extended Data

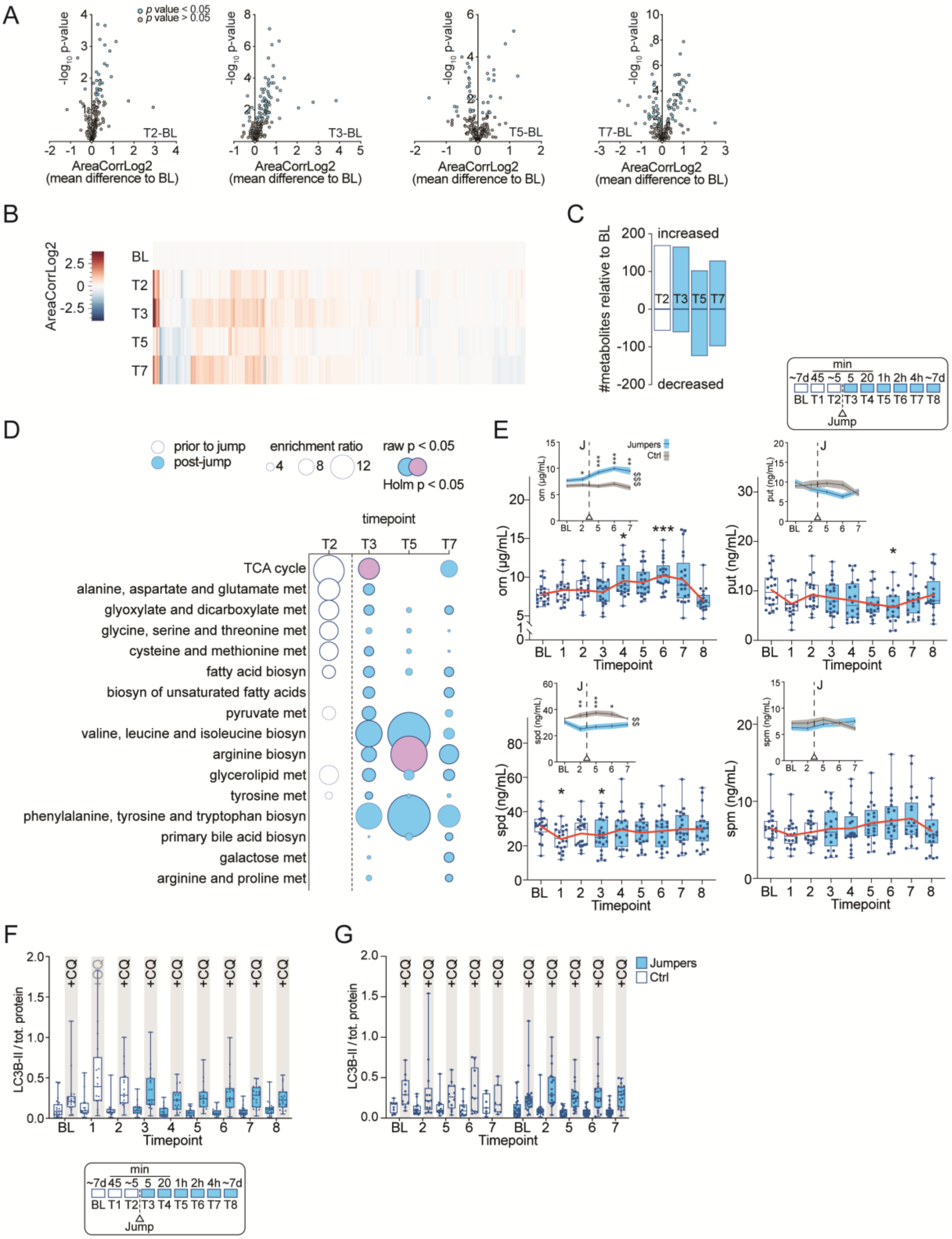

**Extended Data Fig. 1 Changes in metabolites and autophagic flux in bungee jumpers and controls.**

**(A)** Volcano plots of metabolites compared to baseline (BL) within the group of bungee jumpers (cohort #1, *Mixed effects model followed a two-stage linear step-up procedure of Benjamini, Krieger, and Yekutieli to correct for multiple comparisons by controlling the false discovery rate (<0.05); comparisons to T0; n = 22)*.

**(B)** Heatmap of metabolite intensities (AreaCorrLog2Cen) relative to BL in the group of bungee jumpers *(mean difference to BL)*.

**(C)** Number of changed metabolites in response to acute stress relative to BL at specific time points in the bungee jumpers *(FC to BL)*.

**(D)** Metabolite set enrichment analysis based on KEGG human metabolic pathways in the bungee jumpers.

**(E)** Quantitative analysis of orn and polyamine levels within the bungee jumpers and controls over the whole-time course of the experiment *(n = 21 Jumpers, n = 16 Ctrl; orn: Tukey's multiple comparisons tests: Jumpers: BL vs T4: 95% CI: -3259 to -171.4, p = 0.0233; Jumpers: BL vs T6: 95% CI: -3811 to -1098, p = <. 001; Jumpers vs Ctrl: Sidak's multiple comparisons tests: T2: 95% CI:* -2643 to -144,1, *p = 0.0228; T5: 95% CI: -4066 to -1157, p = <. 001: T6: 95% CI: -4481 to -1400, p = <0,001: T7: 95% CI: -5331 to -1036, p = 0,0016. put: Tukey's multiple comparisons test: BL vs. T6: 95% CI: 0.1151 to 6.781, p = 0,0396. spd: Tukey's multiple comparisons tests: BL vs. T1: 95% CI: 0.8586 to 14.06, p = 0,0201; BL vs. T3: 95%CI: 0.09574 to 10.64, p = 0,0441. Jumpers vs. Ctrl: Sidak's multiple comparisons tests: T2: 95% CI: 2.728 to 15.67, p =0,0025; T5 =95% CI: 3.773 to 17.38, p = <.001; T6 = 95% CI: 1.225 to 16.82, p = 0,0171).*

**(F)** LC3B-II levels normalized to total protein in PBMCs from bungee jumpers, determined by immunoblotting. The figure displays the results of LC3B-II levels in blood samples treated with CQ compared to those treated with the veh control.

**(G)** LC3B-II levels normalized to total protein in PBMCs from both bungee jumpers and controls, determined by immunoblotting at selected time points. The representation is similar to that in panel **(F)**.

p ≤ 0.05 (*), p ≤ 0.01 (**), p ≤ 0.001 (***)

**Abbreviations in alphabetical order:** ala, asp, glu met = alanine, aspartate and glutamate metabolism, Aminoacyl-tRNA bio = aminoacyl-tRNA biosynthesis, arg bio = arginine biosynthesis, ARG1 = arginase 1, BL = baseline, but met = butanoate metabolism, β-ala met = β-alanine metabolism, CI = confidence interval, CQ = chloroquine, Ctrl = control, cys and meth met = cysteine and methionine metabolism, D-arg and D-orn met = D-arginine and D-ornithine metabolism, FA bio = fatty acid biosynthesis, FC = fold change, gal met = galactose metabolism, glyox, dicarbox met = glyoxylate and dicarboxylate metabolism, Prim BA bio = primary bile acid biosynthesis, prop met = propanoate metabolism, pur met = purine metabolism, put = putrescine, pyr met = pyruvate metabolism, orn = ornithine, spd = spermidine, spm = spermine, TCA cycle = citrate cycle, uFA bio = biosynthesis of unsaturated fatty acids, val, leu, isoleu bio = valine, leucine and isoleucine biosynthesis.

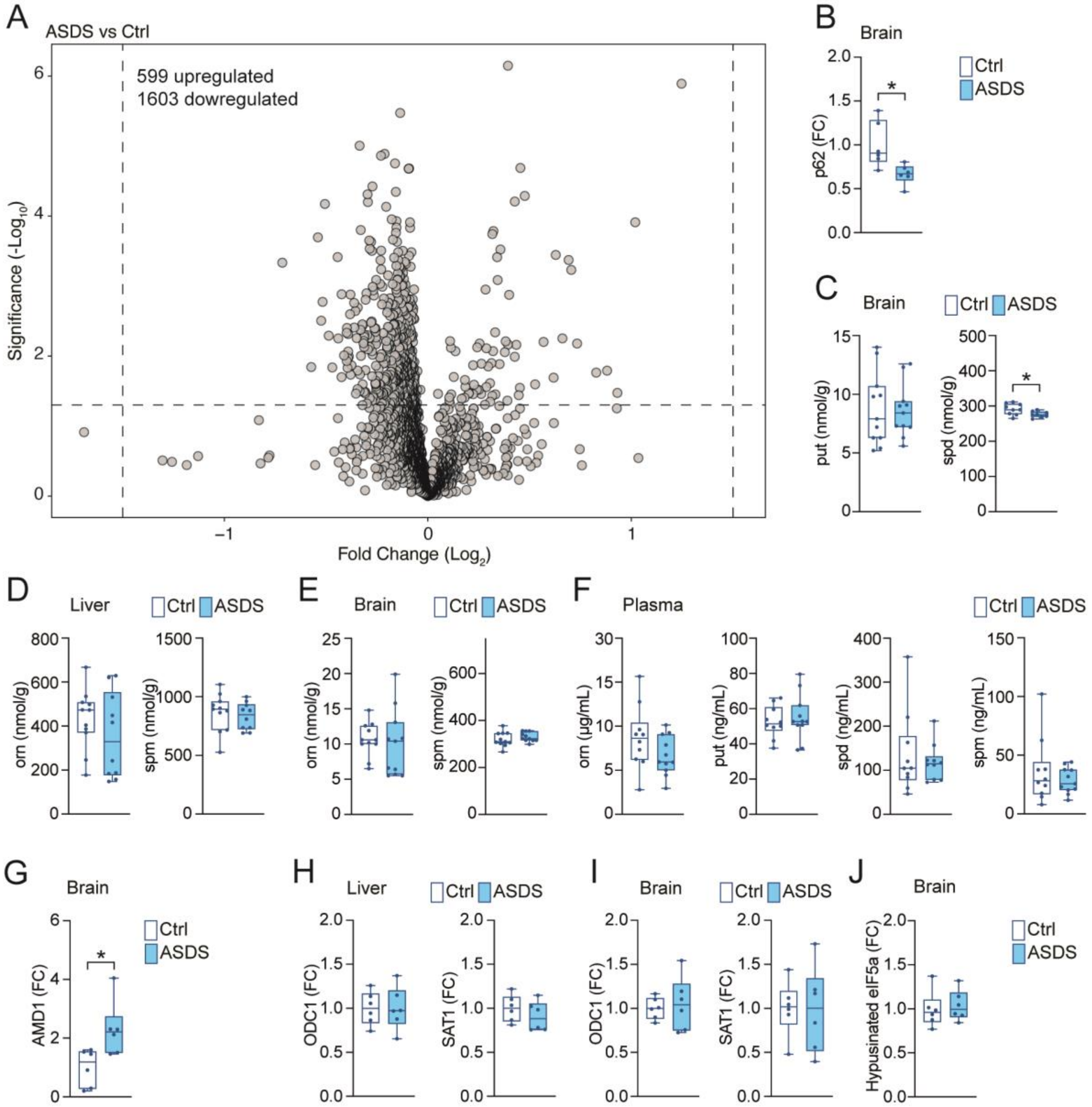

A
ASDS vs Ctrl
599 upregulated
1603 dowregulated
Significance (-Log10)
Fold Change (Log2)
B
Brain
p62 (FC)
Ctrl
ASDS
C
Brain
put (nmol/g)
spd (nmol/g)
D
Liver
orn (nmol/g)
spm (nmol/g)
E
Brain
orn (nmol/g)
spm (nmol/g)
F
Plasma
orn (µg/mL)
put (ng/mL)
spd (ng/mL)
spm (ng/mL)
G
Brain
AMD1 (FC)
H
Liver
ODC1 (FC)
SAT1 (FC)
I
Brain
ODC1 (FC)
SAT1 (FC)
J
Brain
Hypusinated eIF5a (FC)

**Extended Data Fig. 2 Polyamine and protein levels after acute stress in mice.**

**(A)** Volcano plot depicting differentially enriched and depleted proteins in the brains of mice subjected to ASDS compared to control mice. The cutoffs for significant changes are a fold change of ±1.50 and $p < 0.05$ *(mouse #1: n = 6 ASDS, n = 6 Ctrl)*.

**(B)** Quantification of p62 levels in brain lysates from mice in **(A)**, determined by immunoblotting *(n = 6 ASDS, n = 6 Ctrl; two-tailed t-test: p = 0.0165).*

**(C)** Targeted analysis of put and spd levels in brain samples from mice in **(A)** *(spd: n = 8 ASDS, n = 9 Ctrl; unpaired two-tailed t-test: p = 0.0338)*.

**(D)** Targeted analysis of orn and spd levels in liver samples from mice in **(A)**.

**(E)** Targeted analysis of orn and spd levels in brain samples from mice in **(A)**.

**(F)** Targeted analysis of orn, put, spd, and spm levels in plasma samples from mice in **(A)**.

**(G)** Relative AMD1 levels in brain lysates from mice in **(A)**, determined by immunoblotting (*n = 6 ASDS, n = 6 Ctrl; unpaired two-tailed t-test: p = 0.0190).*

**(H)** Relative ODC1 and SAT1 levels in liver lysates from mice in **(A)**, determined by immunoblotting.

**(I)** Relative ODC1 and SAT1 levels in brain lysates from mice in **(A)**, determined by immunoblotting.

**(J)** Relative levels of hypusine normalized to GAPDH in brain lysates from mice in **(A)**, determined by immunoblotting.

$p \leq 0.05$ (*)

**Abbreviations in alphabetical order:** AMD1 = adenosylmethionine decarboxylase 1, ASDS = acute social defeat stress, FC = fold change, GAPDH = glyceraldehyde-3-phosphate dehydrogenase, ODC1 = ornithine decarboxylase 1, orn = ornithine, p62 = SQSTM1 = sequestosome 1, put = putrescine, SAT1 = spermidine/spermine N1-acetyltransferase 1, spd = spermidine, spm = spermine.

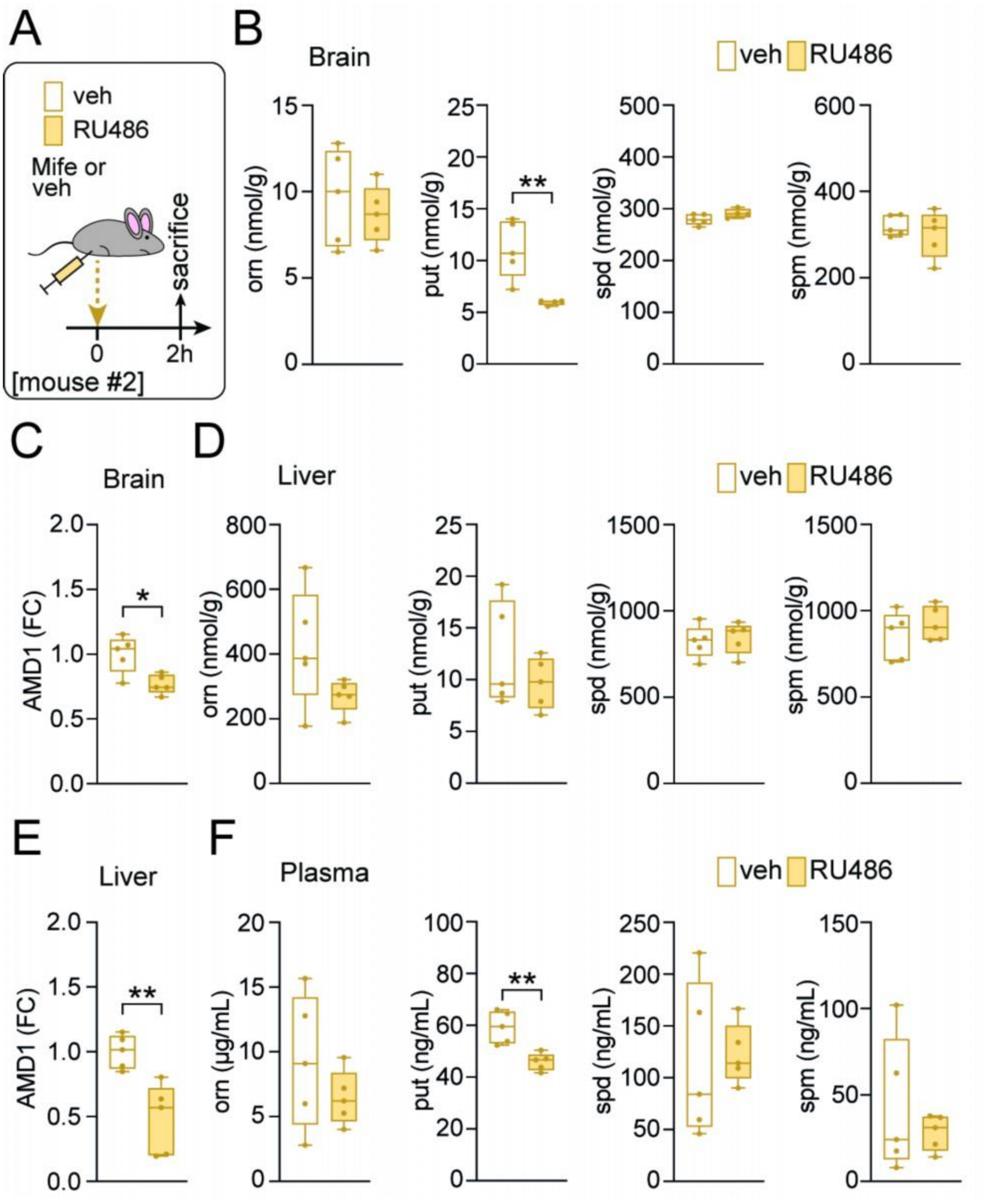
A
veh
RU486
Mife or veh
sacrifice
0
2h
[mouse #2]
B
Brain
veh RU486
orn (nmol/g)
put (nmol/g)
spd (nmol/g)
spm (nmol/g)
**
C
Brain
AMD1 (FC)
*
D
Liver
veh RU486
orn (nmol/g)
put (nmol/g)
spd (nmol/g)
spm (nmol/g)
E
Liver
AMD1 (FC)
**
F
Plasma
veh RU486
orn (µg/mL)
put (ng/mL)
spd (ng/mL)
spm (ng/mL)
**

**Extended Data Fig. 3 Mifepristone injections affect the acute stress response in mice.**

**(A)** Schematic overview of the ASDS paradigm in mice, with mice receiving either the glucocorticoid receptor antagonist RU486 or veh injections *(mouse #2)*.

**(B)** Targeted analysis of orn, put, spd, and spm levels in brain samples from mice in **(A)** *(put: n = 4 RU486, n = 5 veh; unpaired two-tailed t-test: p = 0.0086).*

**(C)** Relative AMD1 levels in brain lysates from mice in **(A)**, determined by immunoblotting *(n = 5 RU486, n = 5 veh; unpaired two-tailed t-test: p = 0.0123).*

**(D)** Targeted analysis of orn, put, spd, and spm levels in liver samples from mice in **(A)**.

**(E)** Relative AMD1 levels in liver lysates from mice in **(A)**, determined by immunoblotting *(n = 5 RU486, n = 5 veh; Mann-Whitney test: p = 0.0048).*

**(F)** Targeted analysis of orn, put, spd, and spm levels in plasma samples from mice in **(A)** *(put: n = 5 RU486, n = 5 veh; unpaired two-tailed t-test: p = 0.0028).*

$p \leq 0.05$ (*), $p \leq 0.01$ (**)

**Abbreviations in alphabetical order:** AMD1 = adenosylmethionine decarboxylase 1, ASDS = acute social defeat stress, FC = fold change, GR = glucocorticoid receptor, orn = ornithine, put = putrescine, RU486 = Mife = mifepristone, spd = spermidine, spm = spermine.

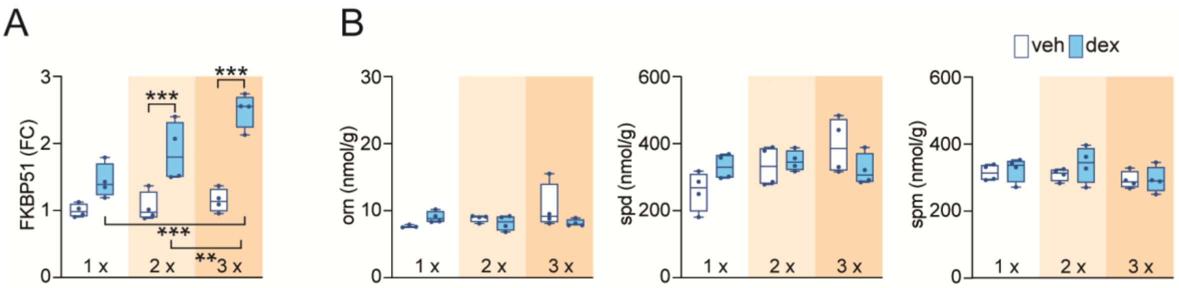


**Extended Data Fig. 4 FKBP51 and polyamine levels in mice subjected to dexamethasone injections.**

**(A)** Relative FKBP51 levels in brain lysates from mice subjected to stepwise dex or veh injections, determined by immunoblotting (*mouse #3, n = 4 dex, n = 4 veh per injection. dex vs. veh: Sidak's multiple comparisons tests: 2x: 95% CI: 0,3343 to 1,312; p = <.001, 3x: 95% CI: 0,8562 to 1,834, p = <.001. Tukey's multiple comparisons tests: Dex: 1x vs. 3x: 95% CI: -0,9061 to 0,04235, p = <.001; 2x vs. 3x: 95% CI: -1,097 to -0,1483, p = 0.009)*.

**(B)** Targeted analysis of orn, spd, and spm levels in brain samples from mice in **(A)**.

p ≤ 0.01 (**), p ≤ 0.001 (***)

**Abbreviations in alphabetical order:** dex = dexamethasone, FC = fold change, orn = FKBP51 = FK506-binding protein 51, ornithine, spd = spermidine, spm = spermine, veh = vehicle.

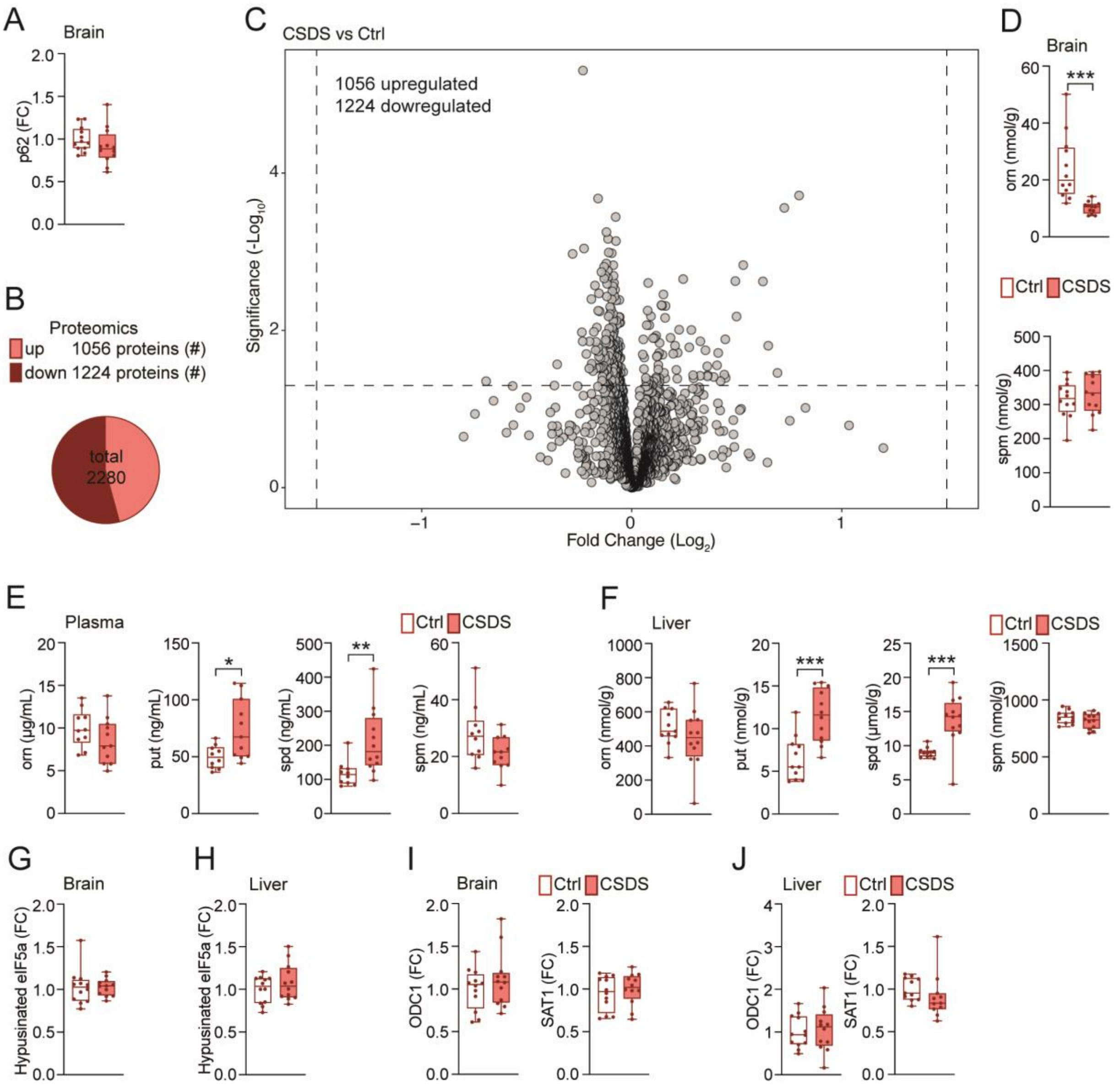

A
Brain
p62 (FC)
B
Proteomics
up 1056 proteins (#)
down 1224 proteins (#)
total 2280
C
CSDS vs Ctrl
1056 upregulated
1224 dowregulated
Significance (-Log10)
Fold Change (Log2)
D
Brain
orn (nmol/g)
***
Ctrl
CSDS
spm (nmol/g)
E
Plasma
orn (µg/mL)
put (ng/mL)
spd (ng/mL)
spm (ng/mL)
*
**
F
Liver
orn (nmol/g)
put (nmol/g)
spd (µmol/g)
spm (nmol/g)
***
G
Brain
Hypusinated eIF5a (FC)
H
Liver
I
Brain
ODC1 (FC)
SAT1 (FC)
J
Liver

**Extended Data Fig. 5 Protein and polyamine analyses in various samples from mice subjected to chronic social defeat stress.**

**(A)** Relative p62 levels in brain lysates from mice subjected to CSDS and controls, determined by immunoblotting *(mouse #4)*.

**(B)** Pie chart illustrating the differential expression of proteins in the brain in response to CSDS compared to control from mice in **(A)**.

**(C)** Volcano plot depicting differentially enriched and depleted proteins in brain samples from mice in **(A)**. The cutoffs for significant changes are FC of ±1.50 and $p < 0.05$ *(n = 12 CSDS, n = 12 Ctrl)*.

**(D)** Targeted analysis of orn and spm levels in brain samples from mice in **(A)** *(orn: n = 12 CSDS, n = 12 Ctrl; unpaired two-tailed t-test: p <.001).*

**(E)** Targeted analysis of orn, put, spd, and spm levels in plasma samples from mice in **(A)** *(put: n = 11 CSDS, n = 10 Ctrl; unpaired two-tailed t-test: p = 0.0131. spd: n = 12 CSDS, n = 10 Ctrl, unpaired two-tailed t-test, p = 0.0081).*

**(F)** Targeted analysis of orn, put, spd and spm levels in liver samples from mice in **(A)** *(n = 12 CSDS, n = 11 Ctrl; unpaired two-tailed t-tests: put: p <.001, spd: p <.001).*

**(G)** Relative levels of hypusine normalized to GAPDH in brain lysates from mice in **(A)**, determined by immunoblotting.

**(H)** Relative levels of hypusine normalized to GAPDH in liver lysates from mice in **(A)**, determined by immunoblotting.

**(I)** Relative ODC1 and SAT1 levels in brain lysates from mice in **(A)**, determined by immunoblotting.

**(J)** Relative ODC1 and SAT1 levels in liver lysates from mice in **(A)**, determined by immunoblotting.

$p \leq 0.05$ (*), $p \leq 0.01$ (**), $p \leq 0.001$ (***)

**Abbreviations in alphabetical order:** Ctrl = control, CSDS = chronic social defeat stress, FC = fold change, GAPDH = glyceraldehyde-3-phosphate dehydrogenase, ODC1 = ornithine decarboxylase 1, orn = ornithine, p62 = SQSTM1 = sequestosome 1, put = putrescine, SAT1 = spermidine/spermine N1-acetyltransferase 1, spd = spermidine, spm = spermine.

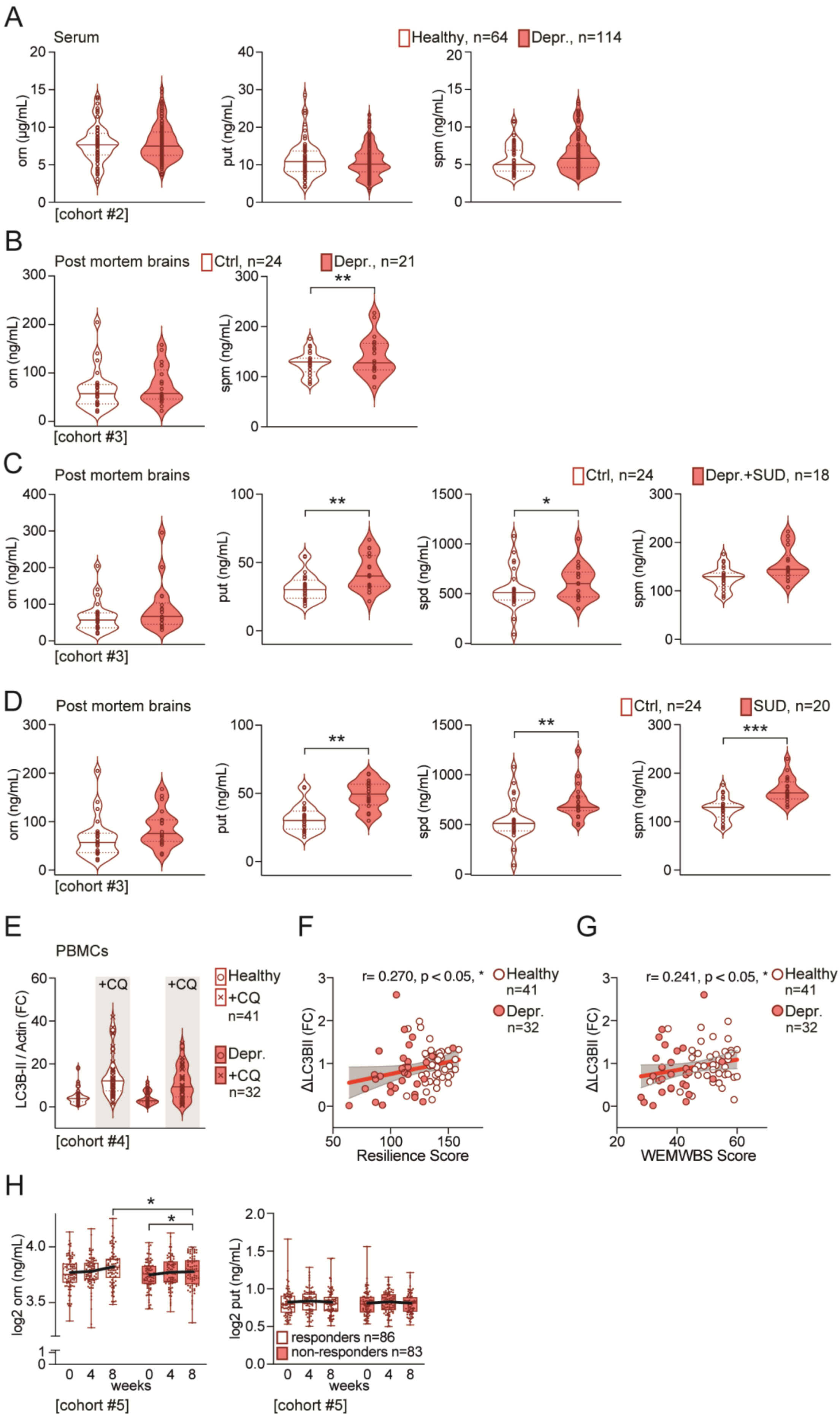
A
Serum
Healthy, n=64
Depr., n=114
orn (µg/mL)
put (ng/mL)
spm (ng/mL)
[cohort #2]
B
Post mortem brains
Ctrl, n=24
Depr., n=21
orn (ng/mL)
spm (ng/mL)
**
[cohort #3]
C
Post mortem brains
Ctrl, n=24
Depr.+SUD, n=18
orn (ng/mL)
put (ng/mL)
spd (ng/mL)
spm (ng/mL)
**
*
[cohort #3]
D
Post mortem brains
Ctrl, n=24
SUD, n=20
orn (ng/mL)
put (ng/mL)
spd (ng/mL)
spm (ng/mL)
**
**
***
[cohort #3]
E
PBMCs
LC3B-II / Actin (FC)
+CQ
+CQ
Healthy
+CQ
n=41
Depr.
+CQ
n=32
[cohort #4]
F
r= 0.270, p < 0.05, *
Healthy
n=41
Depr.
n=32
ΔLC3BII (FC)
Resilience Score
G
r= 0.241, p < 0.05, *
Healthy
n=41
Depr.
n=32
ΔLC3BII (FC)
WEMWBS Score
H
log2 orn (ng/mL)
*
*
weeks
[cohort #5]
log2 put (ng/mL)
responders n=86
non-responders n=83
weeks
[cohort #5]

**Extended Data Fig. 6 Polyamine levels and autophagic flux in depression and substance use disorder.**

**(A)** Targeted analysis of orn, put and spm levels in serum samples from depressed patients and healthy controls in cohort #2 *(n = 114 Depr., n = 64 Healthy)*.

**(B)** Targeted analysis of orn and spm levels in postmortem hippocampal brain samples from subjects with depression and control brain donors from cohort #3 *(n = 21 Depr., n = 24 Ctrl, p-values are derived from ANCOVA reflecting significant effects of age and ZT time of death for orn and presence ethanol in the blood at death and duration of depression for spm).*

**(C)** Targeted analysis of orn, put, spd, and spm levels in postmortem hippocampal brain samples from subjects with depression and SUD, as well as control brain donors from cohort #3 *(n = 18 Depr./SUD, n = 24 Ctrl, p-values are derived from ANCOVA reflecting significant effects of ZT time of death and suicide for orn, age and opioids in the blood at death for spd, presence of SSRIs in the blood at death and ethanol in the blood at death for spm).*

**(D)** Targeted analysis of orn, put, spd, and spm levels in postmortem hippocampal brain samples from subjects with SUD and control brain donors from cohort #3 *(n = 20 SUD, n = 24 Ctrl, p-values are derived from ANCOVA reflecting significant effects of pH for orn, presence of ethanol in the blood at death for put, and PMI for spm).*

**(E)** Relative LC3B-II levels in PBMCs from depresses patients and healthy controls at baseline of cohort #4. Blood samples were either treated with CQ or veh.

**(F)** Correlation analysis between autophagic flux and Resilience Score in depressed patients and healthy controls at baseline measurement of cohort #4 (*Spearman's rank correlation: r = 0.2687, p = 0.025*).

**(G)** Correlation analysis between autophagic flux and WEMWBS Scale in depressed patients and healthy controls at baseline measurement of cohort #4 (*Spearman's rank correlation, r = 0.2417, p = 0.044*).

**(H)** Targeted analysis of orn and put levels in depressed patients from cohort #5 ((responders (n = 86) and non-responders (n = 83)), with measurements taken at baseline (BL), after 4 weeks (W4), and after 8 weeks (W8) of antidepressant treatment. The cohort is divided into responders and non-responders according to changes in HAM-D 17 from BL to W4, with the results highlighting the comparison of polyamine levels between these groups.

$p \leq 0.05$ (*), $p \leq 0.01$ (**), $p \leq 0.001$ (***)

**Abbreviations in alphabetical order:** BL = baseline, Ctrl = control, CQ = chloroquine, Depr. = depressive patients, HAM-D 17 = 17-item Hamilton Depression Rating Scale, LC3B-II = microtubule-associated protein 1 light chain 3 beta-II, PBMC = peripheral blood mononuclear cells, orn = ornithine, put = putrescine, spd = spermidine, spm = spermine, SUD = substance use disorder, veh = vehicle, WEMWBS = Warwick-Edinburgh Mental Well-Being Scale.

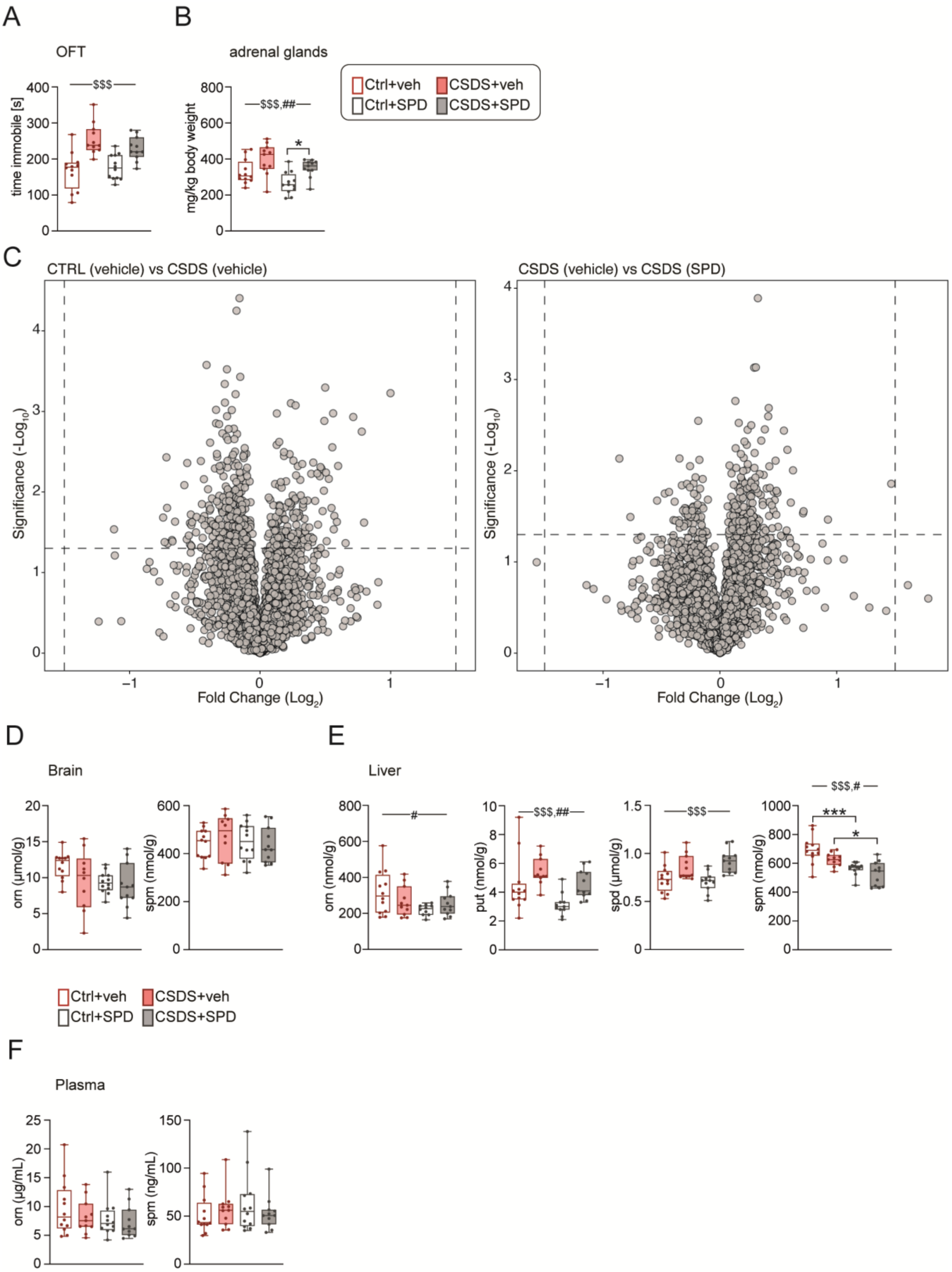

A
OFT
time immobile [s]
$$$
B
adrenal glands
mg/kg body weight
$$$,##
*
Ctrl+veh
CSDS+veh
Ctrl+SPD
CSDS+SPD
C
CTRL (vehicle) vs CSDS (vehicle)
CSDS (vehicle) vs CSDS (SPD)
Significance (-Log10)
Fold Change (Log2)
D
Brain
orn (µmol/g)
spm (nmol/g)
E
Liver
orn (nmol/g)
put (nmol/g)
spd (µmol/g)
spm (nmol/g)
#
$$$,##
$$$
$$$,#
***
*
Ctrl+veh
CSDS+veh
Ctrl+SPD
CSDS+SPD
F
Plasma
orn (µg/mL)
spm (ng/mL)

**Extended Data Fig. 7 Effects of SPD treatment and CSDS on behavior and polyamine levels.**

**(A)** Time spent immobile was measured in an open field test (OFT) in mice divided into four groups: n = 12 Ctrl+veh, n = 12 Ctrl+SPD, n = 11 CSDS+SPD, n = 11 CSDS+veh *(mouse #5: Two-way ANOVA, main CSDS effect:* $F_{(1, 42)} = 31.98$; $p < 0.001$*)*.

**(B)** Normalized adrenal gland weight in mice from **(A)** *(n = 12 Ctrl-veh, n = 12 Ctrl-SPD, n = 11 CSDS+SPD, n = 11 CSDS+veh; two-way ANOVA, main CSDS effect:* $F_{(1, 42)} = 15.40$, $p < 0.001$*, main SPD effect:* $F_{(1, 42)} = 7.970$, $p = 0.0072$*. Tukey's multiple comparisons test for Ctrl+SPD vs. CSDS+SPD: 95% CI: -0.07967 to -0.004796, p = 0.022).*

**(C)** Volcano plots illustrating differentially expressed proteins in the brains of mice subjected to CSDS compared to controls, as well as those i.p. injected with SPD versus those treated with veh in the CSDS model. The cutoffs for significant changes are fold change of ±1.50 and p < 0.05 *(n = 8 Ctrl+veh, n = 8 Ctrl+SPD, n = 8 CSDS+SPD, n = 7 CSDS+veh).*

**(D)** Targeted analysis of orn and spm levels in brain samples from mice in **(A)**.

**(E)** Targeted analysis of orn, put, spd, and spm levels in liver samples from mice in **(A)** *(n = 12 Ctrl+veh, n = 10 Ctrl+SPD, n = 11 CSDS+SPD, n = 11 CSDS-veh; orn: main SPD effect: Two-way ANOVA:* $F_{(1, 40)} = 4.539$, $p = 0.0393$*; put: Two-way ANOVA: main CSDS effect:* $F_{(1, 41)} = 15.78$, $p < .001$*, main SPD effect:* $F_{(1, 41)} = 9.574$, $p = .004$*. spd: Two-way ANOVA: main CSDS effect:* $F_{(1, 41)} = 22.25$, $p < .001$*. spm: Two-way ANOVA: main CSDS effect:* $F_{(1, 41)} = 6.395$, $p = 0.015$*, main SPD effect:* $F_{(1, 41)} = 29.42$, $p < .001$*. Tukey's multiple comparisons test: Ctrl+veh vs. Ctrl+SPD: 95% CI: 51.59 to 212.8, p = < .001; CSDS+veh vs. CSDS+SPD: 95% CI: 18.88 to 183.6, p = 0.011).*

**(F)** Targeted analysis of orn and spm levels in plasma samples from mice in **(A)**.

p ≤ 0.05 (*), p ≤ 0.01 (**), p ≤ 0.001 (***)

**Abbreviations in alphabetical order:** Ctrl = control, CSDS = chronic social defeat stress, orn = ornithine, OFT = open field test, put = putrescine, SPD = spermidine i.p., spd = spermidine (metabolite), spm = spermine, $ = CSDS effect, # = SPD effect.

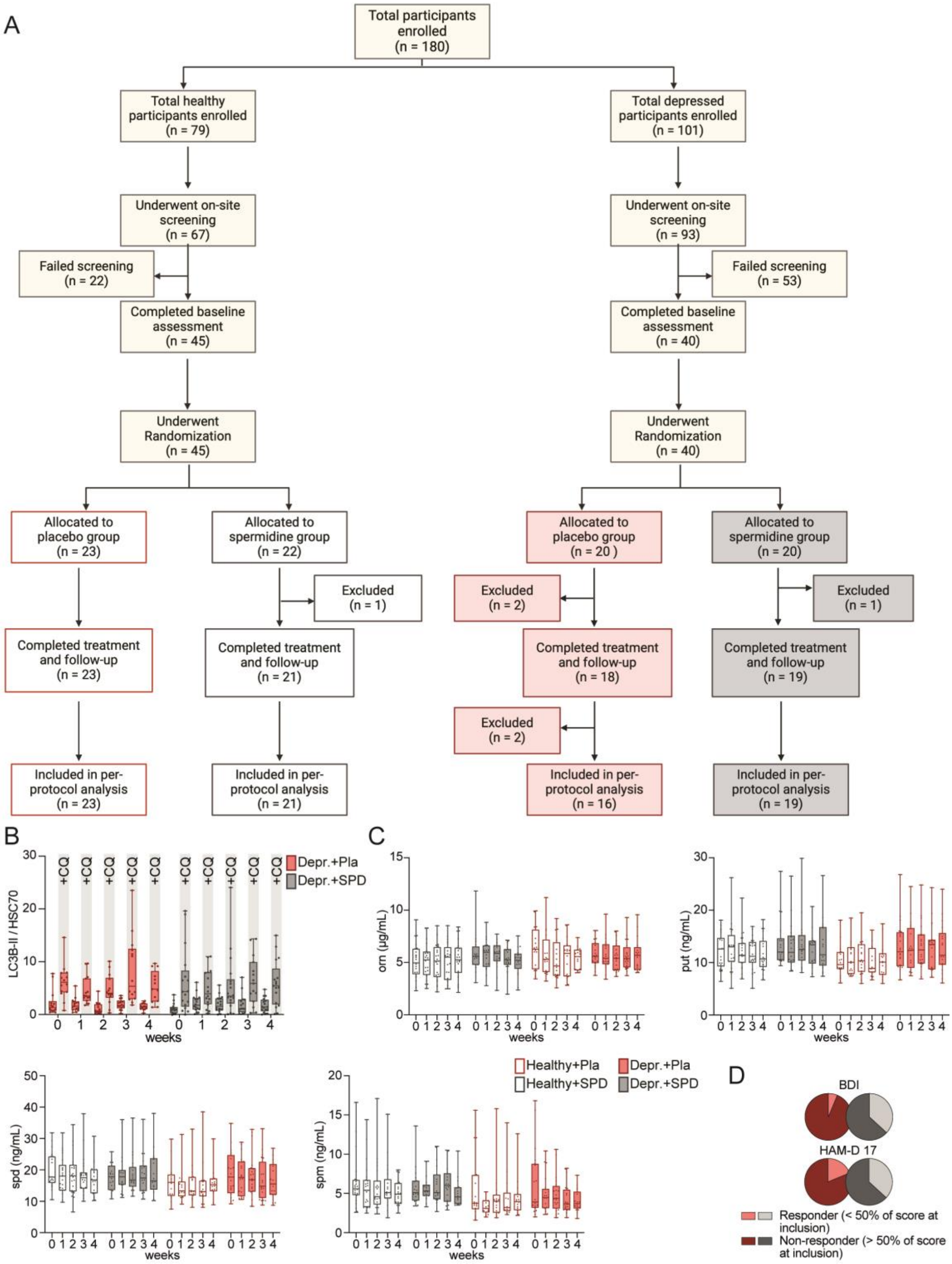
A
Total participants enrolled (n = 180)
Total healthy participants enrolled (n = 79)
Total depressed participants enrolled (n = 101)
Underwent on-site screening (n = 67)
Underwent on-site screening (n = 93)
Failed screening (n = 22)
Failed screening (n = 53)
Completed baseline assessment (n = 45)
Completed baseline assessment (n = 40)
Underwent Randomization (n = 45)
Underwent Randomization (n = 40)
Allocated to placebo group (n = 23)
Allocated to spermidine group (n = 22)
Allocated to placebo group (n = 20 )
Allocated to spermidine group (n = 20)
Excluded (n = 1)
Excluded (n = 2)
Excluded (n = 1)
Completed treatment and follow-up (n = 23)
Completed treatment and follow-up (n = 21)
Completed treatment and follow-up (n = 18)
Completed treatment and follow-up (n = 19)
Excluded (n = 2)
Included in per-protocol analysis (n = 23)
Included in per-protocol analysis (n = 21)
Included in per-protocol analysis (n = 16)
Included in per-protocol analysis (n = 19)
B
LC3B-II / HSC70
+CQ
Depr.+Pla
Depr.+SPD
weeks
C
orn (µg/mL)
put (ng/mL)
spd (ng/mL)
spm (ng/mL)
weeks
Healthy+Pla
Healthy+SPD
Depr.+Pla
Depr.+SPD
D
BDI
HAM-D 17
Responder (< 50% of score at inclusion)
Non-responder (> 50% of score at inclusion)

**Extended Data Fig. 8 Analysis of SPD treatment effects and response in depressed patients.**

(**A**) Study flowchart of cohort #4. The per-protocol set included all participants who completed the 3-week intervention and follow-up appointment and were not excluded based on the exclusion criteria.

**(B)** LC3B-II levels normalized to HSC70 in PBMCs from depressed patients who received SPD and those who received Pla in cohort #4. Blood samples were either treated with CQ or veh.

**(C)** Targeted analyses of orn, put, spd, and spm levels in plasma samples from cohort #4.

**(D)** Pie charts showing responder and non-responder status in BDI and HAM-D 17 from cohort #4. Responders are characterized by at least 50% reduction from the baseline score; non-responders are defined as less than 50% reduction from baseline score *(BDI: responder: SPD: 7 out of 19, Pla: 1 out of 16, non-responder: SPD: 12 out of 19, Pla: 15 out of 16, HAM-D 17: responder: SPD: 7 out of 19, Pla: 3 out of 16, non-responder: SPD: 12 out of 19, Pla: 13 out of 16).*

**Abbreviations in alphabetical order:** BDI = Beck Depression Inventory, CQ = chloroquine, Depr. = depressive patients, HAM-D 17 = 17-item Hamilton Depression Rating Scale, HSC70 = heat shock cognate, orn = ornithine, Pla = placebo, put = putrescine, SPD = spermidine (supplement), spd = spermidine (metabolite), spm = spermine.

### Supplementary Information

**Supplementary Table 1** – Demographic information of bungee jump participants and healthy control group from cohort #1.

**Supplementary Table 2** – Metabolomic profile of cohort #1, measured in plasma samples using high-resolution mass spectrometry.

**Supplementary Table 3** – Proteomic measurements of whole-brain extracts of mice after ASDS and controls.

**Supplementary Table 4** – Proteomic measurements of whole-brain extracts of mice after CSDS and controls.

**Supplementary Table 5** – Demographic information of depressed and healthy participants from cohort #2.

**Supplementary Table 6** – Demographic information of brain donors who provided postmortem samples for cohort #3.

**Supplementary Table 7** – Demographic information of depressed and healthy participants from cohort #4.

**Supplementary Table 8** – Demographic information of depressed participants from cohort #5.

**Supplementary Table 9** – Proteomic measurements of whole-brain extracts of mice after CSDS, with either SPD or veh injections, and control groups.

### Source Data Table

Table includes raw data as presented for all main and extended data figures including uncropped Western blot images and raw files of metabolomic measurements.

### Materials and Methods

#### Mouse

In this manuscript, we present the results of various mouse experiments. The following section describes each experiment, including the location where it was conducted. Next, we outline the sample preparation, which was identical for most experiments. Any variations in the subsequent analyses are indicated in the text. All experiments were conducted in compliance with relevant ethical regulations, as detailed in the respective sections.

#### Mouse #1, #2, #4

Mouse experiments were approved by the local authorities from Rhineland Palatinate (Animal Protection Committee of the State Government, Landesuntersuchungsamt Rheinland-Pfalz, Koblenz, Germany) and conducted in accordance with the European Communities Council Directive 86/609/EEC. Single-housed adult male C57BL/6J (B6J) mice, 8 to 10 weeks old, were provided with *ad libitum* access to food and water. Sample collection was performed in collaboration between the University Medical Center Mainz, Germany and the Neurohomeostasis Research Group in Bonn, Germany. Following steps were performed by the Neurohomeostasis Research Group in Bonn, Germany and at the Institute of Molecular Biosciences at the University of Graz, Austria.

#### Social defeat stress protocol

In the ASDS experiment (mouse #1), a total of 12 B6J mice were utilized and divided into two groups: an experimental group (n = 6) that was exposed to an aggressor mouse for 5 to 10 min, and a control group (n = 6).

In a subsequent experiment (mouse #2) the ASDS design was employed again, involving a total of 24 B6J mice divided into four groups: two experimental groups subjected to acute social defeat stress and receiving either glucocorticoid receptor antagonist RU486 (n = 6) or corn oil (n = 6) injections, and two control groups receiving either glucocorticoid receptor antagonist RU486 (n = 6) or corn oil (n = 6) injections.

In a third experiment (mouse #4), animals were subjected to CSDS as previously described in[1]. The CSDS was administered over 10 days, during which the experimental group (n = 12) was subjected to daily defeat sessions. Control mice (n = 12) were placed in a novel cage for 90 s over 10 consecutive days, and their cages were also equipped with a metal grid to mimic the conditions experienced by the defeated mice.

Retroorbital blood sampling was performed under terminal anesthesia followed by the sacrifice of the mice. Tissues were then promptly dissected and snap-frozen in liquid nitrogen. For this manuscript, the brain and liver samples were analyzed.

**Mouse #3**

Mouse experiments, behavioral tests, and sample collection were performed at the Max Planck Institute in Munich, Germany. Following steps were performed by the Neurohomeostasis Research Group in Bonn, Germany and at the Institute of Molecular Biosciences at the University of Graz, Austria. This animal study was conduced in accordance with the recommendations of the Federation for Laboratory Animal Science Associations and was approved by the Government of Upper Bavaria (AZ: 55.2-1-54-2532, Vet 02-17-206).

**Animals and animal housing**

Mice were bred at the Max Planck Institute of Biochemistry, Martinsried-Planegg, Germany and housed at the facilities of the Max Planck Institute of Psychiatry, Munich, Germany under standard housing conditions (23 ± 3 °C, 40 ± 10% relative humidity). Group-housed adult male C57BL/6N (B6N) mice, approximately 2 to 3 months old, were provided with ad libitum access to food (Altromin Haltungsdiät 1328 or Zuchtdiät, Altromin Spezialfutter, Lage, Germany) and water. They were kept in IVC cages (Greenline, Tecniplast) equipped with bedding, nesting material, and enrichment items, including a wooden rodent tunnel (4.5 × 4 cm, diameter: 30 mm, ABEDD, Vienna, Austria).

**Dex injections**

Experiments were conducted during the mice's active phase (lights off). For the procedures, 20 mg/kg of the substance was dissolved in saline (NaCl) for intraperitoneal (i.p.) injections. The dex administration involved six experimental groups (n = 24). The experimental groups received either a single dex injection (20 mg/kg i.p.), two dex injections spaced four hours apart, or three dex injections, each spaced four hours apart. Control groups followed the same schedule but received vehicle injections instead. After administering the treatments, mice were euthanized using an overdose of isoflurane, followed by organ collection and final blood sampling. For this manuscript, brain samples were analyzed.

**Mouse #5**

Mouse experiments, behavioral tests, and sample collection were performed in the animal facility of Mailman Research Center at McLean Hospital in Belmont, Massachusetts, USA. Afterwards material was stored at -80 °C and shipped on dry ice to the Research Group Neurohomeostasis in Bonn, Germany, where further sample preparation steps and analysis were performed. Additionally, mass spectrometric analyses were conducted at the Institute of Molecular Biosciences at the University of Graz, Austria. This animal study was approved by the McLean Hospital Institutional Animal Care and Use Committee and complied with the National Institutes of Health guidelines.

**Animals and animal housing**

Eight-week-old male B6J mice (n = 47) were purchased from The Jackson Laboratory (Bar Harbor, ME). Twelve-week-old CFW male mice (n = 30) were pair-housed with ovariectomized (OVX) CFWs. Eight-week-old group-housed CFW males (n = 5/cage; n = 30) served as social stimulus mice during pre-defeat aggression tests and open field social interaction tests. Group-housed OVX CFW mice (n = 24) served as stimulus mice for home cage social interaction tests. CFW mice were purchased from Charles River Laboratories (Wilmington, MA). All animals were housed under standard laboratory conditions with a 12 h light–dark cycle (lights on from 07:00 to 19:00) in clear Plexiglas cages (19 x 29 x 13 cm) with unrestricted access to food (Purina Laboratory Rodent Diet 5001) and water.

**I.p. SPD treatment**

For four weeks, male B6J mice received either daily i.p. injections of 10 mg/kg spermidine (SPD, Sigma-Aldrich, St. Louis, Missouri, #S2626) or saline vehicle at an injection volume of 10µL/g body weight.

**Twenty-one-day CSDS**

The 21 day CSDS paradigm was performed as described previously[2]. Twelve-week-old CFW male mice were pair-housed with ovariectomized CFWs to foster territorial home cage aggression. Resident-intruder confrontations were conducted on three consecutive days[3]; briefly, the OVX mouse was removed from the resident home cage and replaced with a group-housed male CFW intruder. Highly aggressive resident mice-initiated attacks toward the intruder within 30 seconds (n = 26/30); these aggressive residents were single-housed and used to defeat experimental B6J male mice during the 21 day CSDS protocol. On defeat days, experimental B6J mice were introduced into the home cage of an unfamiliar aggressive resident for a single bout of aggression (<1 min). After the defeat, experimental mice were separated from the aggressor by a perforated Plexiglas divider, preventing physical but allowing sensory contact for 24 h. Each day, experimental B6J mice were defeated and housed adjacent to an unfamiliar aggressive CFW resident. Non-defeated control B6J male mice were housed adjacent to another control in their home cages for the 21 day CSDS. Experimental and control B6J mice were handled daily during i.p. saline or SPD injections. Injections always took place immediately before the daily defeat sessions. Behavioral tests were conducted in the third week of CSDS prior to daily defeats. All mice were sacrificed in the morning the day after the last defeat.

**Behavior**

Experiments were analyzed using the automated video-tracking system ANYmaze (Stoelting, Wood Dale, IL), unless stated otherwise.

**Open field**

Testing was performed in an open field arena (50 x 50 x 50 cm) dimly illuminated (10 lux). All mice were placed into a corner of the apparatus at the beginning of the trial. The testing duration was 10 min and the distance traveled and the time immobile was assessed.

**Social avoidance test**

The social avoidance test was performed as described previously[4]. B6J mice explored an open field arena (50 x 50 x 50 cm) containing an empty wire cup (10 cm diameter) for a 2.5-min non-target trial. Then, an unfamiliar non-aggressive male CFW mouse was placed in the wire cup and B6J mice explored the arena and the caged CFW mouse during a 2.5-min target trial. Distance traveled during non-target and the target trials were assessed. To quantify social approach behavior, an interaction zone was defined as the area surrounding the wire cup. Entries into and time spent in the interaction zone were quantified during non-target and target trials; social interaction ratios were calculated as entries into or time in the interaction zone during the target trial/non-target trial.

**Home cage social interaction test**

The home cage social interaction test was performed as described previously[5]. Briefly, social contact with a non-aggressive, group-housed ovariectomized CFW female stimulus mouse was evaluated during 1.5-min tests conducted in a fresh cage. Tests were video recorded (monochromatic recordings with a frame rate of 30) using an overhead Logitech C922 Pro Stream webcam. Training frames (n = 214) were annotated in multi-animal DeepLabCut (maDLC)[6]; videos were analyzed in maDLC to yield pose estimations that served as the input for Simple Behavioral Analysis (SimBA)[7]. Frames were labelled for random forest classification of approach (n = 1481), social contact (n = 2001), avoidance (n = 1076), freezing (n = 2162), and escape (n = 573). These classifiers were operationally defined as: *approach* as diminishing distance between B6J and OVX mice while the B6J mouse is traversing toward the OVX mouse; *social contact* as physical contact with the OVX mouse, initiated by the B6J mouse; *freezing* as motionless B6J during bouts of contact initiated by the OVX mouse; and *avoidance* as B6J orientation toward the OVX mouse while maintaining at least one body length distance between themselves and the OVX mouse[5,8,9]. Analysis of historical videos revealed $r^2 > 0.8$ for human vs. machine behavioral classification results.

**Sampling procedure**

Mice were sacrificed by decapitation following quick anesthesia by isoflurane. Brains and other organs were removed, snap-frozen in isopentane at -40 °C, and stored at -80 °C. Adrenal glands were removed, dissected from fat and weighed.

**Statistical analysis**

This experiment was conducted as a 2 x 2 between-subjects design with saline + non-defeated, SPD + non-defeated, saline + defeated, and SPD + defeated groups of B6J male mice. Data were analyzed in Graphpad Prism (7.0 or 9.0). For four group comparisons, two-way analyses of variance (ANOVA) were performed, followed by Tukey's post hoc tests ($\alpha$ = 0.05). One mouse from the CSDS vehicle group was excluded from the entire study (including molecular analysis) because it was identified as a significant outlier in multiple behavioral tests based on the Grubbs' outlier test.

**Tissue lysis of mouse brain hemispheres and liver**

Half of the mouse brain hemispheres were lysed using 300 µL of T-PER extraction buffer (Thermo Scientific, 78510), freshly supplemented with 1x cOmplete™ EDTA-free protease inhibitor cocktail (Roche, 4693132001) and 1× PhosSTOP™ phosphatase inhibitor cocktail (Roche, 4906837001) and consequently homogenized using a tissue grinder to produce a smooth lysate. Subsequently, the samples were incubated for 60 min on ice. The lysate was then centrifuged for 10 min at 16,000 x g at 4 °C to separate the supernatant, the latter of which was extracted and added to a new microcentrifuge tube.

The same steps were applied to the liver samples. For this purpose, a piece approximately the size of half a mouse brain hemisphere was excised.

**Samples quantification**

Each sample was quantified using the Bicinchoninic acid assay (BCA Protein Assay Kit, Thermo Scientific, 23225). The aliquots to be quantified were diluted twenty times before adding them to a 96-well microplate (Thermo Scientific, 2205). Afterwards the working solution was prepared by mixing reagent A and reagent B with ratio A:B in a ratio of 50:1. 200 µL of the working solution was then added, and the microplate was incubated at 37 °C for 30 min. Finally, the microplate was quantified using SPARK® Multimode Microplate Reader and SPARKControl Magellan analysis software.

**Sample preparation**

Samples were prepared for separation using an SDS-PAGE system by adding 4× Laemmli sample buffer (Bio-Rad, 1610747) containing 100 mM DTT (Carl Roth, 6908.1) and then heated at 95°C for 5 min.

**Immunoblotting**

After lysis and preparation, protein extracts corresponding to 30 µg protein were loaded onto 10 to 15% SDS-PAGE, and electrophoresis was performed at 80 V for 10 min and then at 150 V for 1 h in running buffer (10% of final concentration 10x Tris/Glycine/SDS Electrophoresis

Buffer, BioRad, 1610772). Proteins were transferred to methanol-activated PVDF membranes (BioRad, 1620177) for 60 min at 350 mA using transfer buffer (10% of final concentration 10x Tris/Glycine/SDS Electrophoresis Buffer, 20% of final concentration methanol, Roth, 4627.5). The membranes were then incubated in Tris-buffered saline, supplemented with 0.05% Tween (Sigma-Aldrich, 9005-64-5), and 5% non-fat milk (Roth, 68514-61-4) for 1 h at room temperature and then incubated with the primary antibody (diluted in TBS/0.05% Tween) overnight at 4 °C while shaking. The following antibodies were used as primary antibodies: AMD1 (1:1000, Abcam, #ab127576), Arginase1 (1:1000 Cell Signaling Technologies, #93668), eIF5A (1:1000, Cell Signaling Technologies, #20765), FKBP5 (1:1000, Cell Signaling, #12210), GAPDH (1:1000, Cell Signaling Technologies, #5174), Hsp70 (1:1000, Cell Signaling Technologies, #4876), Hypusine (1:1000, Millipore, #ABS1064), ODC1 (1:1000, Thermo Fisher Scientific, #PA5-21362), p62 (1:1000, Cell Signaling, #5144S), SAT1 (1:1000, Cell Signaling, #61586), Vinculin (1:1000, Cell Signaling Technologies, #13901).

Secondary antibodies: anti-mouse IgG, HRP-linked antibody (1:10,000, Cell Signaling Technologies, #7076), anti-rabbit IgG, HRP-linked antibody (1:10,000, Cell Signaling Technologies, #7074).

Subsequently, membranes were washed three times with Tris-buffered saline, supplemented with 0.05% Tween, for 10 min and probed with the respective horseradish peroxidase secondary antibody for 1 h at room temperature. The immuno-reactive bands were visualized using Clarity or Clarity Max ECL Western Blotting Substrates (BioRad, 1705060/1705062). Determination of the band intensities were performed with BioRad, ChemiDoc MP Imaging System. Band intensities were quantified using ImageLab 6.0.1 (BioRad) using the rectangular volume tool with local background adjustment.

In general, protein quantification was performed by normalization to the intensity of either GAPDH, Vinculin or Hsp70, which were determined on the same membrane. For the quantification of hypusination, this signal was, when possible, normalized to the signal intensity of the corresponding total protein. Additionally, a specific protein lysate was included in all blots to serve as a reference and facilitate inter-blot comparisons.

**Sample preparation for proteomic analysis**

50 µg lysate was subjected to proteolytic digestion using a modified single-pot solid-phase-enhanced sample preparation (SP3) protocol[10]. Resulting peptides were desalted, dried by vacuum centrifugation and dissolved in 20 µL 0.1% formic acid. A micro-flow LC-MS/MS system composed of a modified Dionex UltiMate 3000 RSLCnano System coupled to a Q Exactive HF-X mass spectrometer (Thermo Fisher Scientific) was used, as described in detail by Bian et al.[11]. Peptides (4.0 µg) were separated on a 15 mm long C18 column with an inner diameter (ID) of 1 mm (Acclaim PepMap RSLC, Thermo Fisher Scientific). A binary 90 or 120-min gradient of water (A) and acetonitrile (B) containing 0.1% (v/v) formic acid and 3% DMSO was applied. Sample loading and column wash was performed at an increased flow rate of 100 µL/min. The column was heated to 55°C. MS1 spectra were acquired at a resolution of 120,000, scan range from 360 to 1400 m/z, a maximum injection time of 100 ms and AGC target of 3E6. Top 20 precursors were subjected to higher-energy c-trap dissociation with a normalized collision energy of 28%.

A resolution of 15,000, AGC target of 1E5, isolation window of 1.6 m/z and a maximum injection time of 22 ms was applied. The dynamic exclusion was set to 30 s.

**Proteomics data processing protocol**

Maxquant software (version 1.6.3.4) was used for data analysisc[12] and searched against a reviewed canonical FASTA database of Mus musculus (UniProt, download: April 11h 2022, 17107 entries). Peptide masses were recalibrated within a window of 20 ppm, the option first search was used. Main search was performed for peptides and peptide fragments within a mass tolerance of 4.5 and 20 ppm respectively. N-terminal acetylation and oxidation of methionine were set as variable, carbamidomethylation of cysteine as static modification. False discovery rate (FDR) was adjusted to less than 1% for both peptides and proteins.

**Proteomic data analysis protocol**

The Perseus software suite (v. 1.6.14.0)[13] was used to filter out protein groups identified only by site, contaminants and reverse hits. Protein groups that were not present in at least 70% of the replicates for any given condition were removed. Log(2) transformation was then applied to the data. To identify significantly regulated protein groups influenced by individual factors or their combination in a bifactorial design, a two-way analysis of variance (ANOVA) was conducted. Significantly regulated proteins were clustered based on their fold change relative to the control group. Additionally, to visualize the top 30 altered protein groups sorted according to p-value under the CSDS condition, k-means clustering was applied, resulting in three distinct clusters, which were then represented in a heatmap using Morpheus (https://software.broadinstitute.org/morpheus). To compare individual groups, Student's t-tests were performed with false discovery rate (FDR) correction to account for multiple testing.

Volcano plot analysis was also carried out by VolcaNoseR[14], using cutoffs of FC ± 1.5 and $p < 0.05$.

#### Plasma and tissue polyamines

Samples were shipped on dry ice to the Institute of Molecular Biosciences at the University of Graz, Austria. Polyamine estimation was performed using LC-MS according to Magnes et al., 2014)[15] with modifications described in[16] and using a modified carbonate buffer (1M ammonium bicarbonate) that was adjusted to pH 10 using hydrochloric acid. Briefly, either 100 µL of plasma was used and processed as described in[15] or frozen tissues (liver, brain) were weighed and appropriate amounts (20 to 80 mg) were extracted with varying volumes of ice-cold 5% trichloroacetic acid (TCA) to yield a final spd concentration of 50 to 1,000 ng/mL (corresponds to the validated linear range of detection) in the final extract prior to polyamine derivatization by isobutyl chloroformate. The extraction solution contained 100 ng/mL isotope-labeled polyamines that served as internal calibration standards and external calibration solutions spiked with the same isotope standard mix were adapted according to the expect range of polyamines for each tissue. Samples were homogenized with an UltraTurrax device for 10 to 15 s followed by incubation on ice for 1 h with vortexing every 15 min. For polyamine analysis of human brain tissue (see below cohort #3), frozen hippocampal tissue slices were scraped off from glass slides using a metal scalpel with continuous cooling on dry ice prior to TCA extraction.

#### Humans

In this manuscript, we present the results of various human cohorts. Serum and plasma analyses were primarily performed on samples from these cohorts. The plasma isolation is described in the section on cohort #1. The plasma and tissue analyses can be found in the section "Mouse". In cohorts #1 and #4, additional analyses were also performed, including PBMC isolation and immunoblots, which are described in the cohort #4 section. All other specific details can be found in the respective sections of each cohort.

#### Cohort #1 - HighStress study

#### Study Design

The aim of this study was to analyze the effects of an acute stress event in the form of a bungee jump on various molecular and neuropsychological parameters. It was a monocenter, non-blinded, non-randomized prospective trial. The intervention group (n = 22) performed a bungee jump, while the control group (n = 16) was not exposed to any experimental stress setup.

Participants were recruited between May 2021 and January 2022. The study was conducted with healthy, male participants.

In the control group, 10 participants completed all designated time points, while the remaining control participants only completed a subset of time points. For the targeted polyamine and flux analyses, all available participants, excluding technical outliers, were included to maximize sample size and statistical power. For all other analyses, only data from the 10 participants who completed all time points were utilized.

**Ethics**

The study was carried out on the premises of the Telekom Baskets Bonn and at the Department of Psychiatry and Psychotherapy, University Hospital Bonn, Germany. The trial protocol was approved by the Ethics Committee of the University Hospital Bonn, Germany (lfd. Nr. 028/21, ClinicalTrials.gov identifier: NCT05144022). The study was conducted in accordance with the Declaration of Helsinki.

**Participants**

After potential participants expressed their interest in performing a bungee jump, they were contacted by the study team to evaluate possible participation. All participants were male and had no previous illnesses. None had ever done a bungee jump before. Exclusion criteria included any neurological or psychiatric pre-existing conditions, regular medication (except for oral contraceptives and thyroid medication), pathological fear of heights, excessive alcohol consumption, smoking, illegal drug use, cardiovascular or pulmonary diseases, serious eye diseases, spine fractures, and surgeries within the last 4 to 6 months. The same inclusion and exclusion criteria applied to the control subjects as to the intervention group.

**Procedures**

**Intervention Group (Bungee Jump Group)**

At the baseline appointments (and at every other blood draw), participants were asked not to eat or drink (except for water) for at least 12 hours. Participants were informed in advance (by email and/or telephone) about the experimental setup. After arriving in the examination room, participants provided written informed consent and underwent further screening assessments for eligibility. This included verbal queries of inclusion and exclusion criteria, physical, neurological, and psychopathological examinations, and an electrocardiogram conducted by medical doctors. If any abnormalities were found, indicating the participant did not meet the inclusion criteria or met the exclusion criteria, they were excluded from the study. Participants also provided a urine sample, tested for illegal substances. If illegal substances were found, the participant was excluded from the study. If no reasons for exclusion were found,

participants completed questionnaires on demographics and mental state, followed by the measurement of pulse and blood pressure and a venous blood sample (T0, baseline).
7 to 10 days after the baseline appointment, the bungee jump took place. Again, all participants were asked not to eat or drink (except for water) for at least 12 hours. In total, seven venous blood draws were taken on the day of the jump. The first blood draw (T1) occurred upon arrival at the experimental site, on average 45 minutes before the jump, followed by the second blood draw on the ground shortly on average 5 min before the jump (T2) and the third blood draw on average 5 min after the jump (T3). Additional blood draws were performed 20 min after the jump (T4), 1 h after the jump (T5), 2 h after the jump (T6), and 4 h after the jump (T7). Participants fasted throughout this period. Between blood samples, participants were asked to avoid movement and stay at the experimental site.

For the bungee jump, participants were lifted on a platform to a height of 60 m by a crane. The technical execution of the jump, including safety measures, was carried out by an external company specializing in bungee jumping. The follow-up appointment (T8) took place 7 to 10 days after the day of the jump. Again, blood was taken for PBMC and plasma isolation, and participants completed the same questionnaires as at T0.

**Control Group**

The control group was not exposed to any experimental stress setup and had only two appointments. On the first appointment (baseline, T0), the same procedures, including neuropsychological testing and blood draw, were conducted as in the intervention group. On the second appointment, venous blood draws were taken at four different time points: upon arrival at the experimental site (T1), 1 h after T1 (T2), 2 h after T1 (T3), and 4 h after T1 (T4). Participants fasted for the entire period. All psychological questionnaires and neuropsychological testing were performed at the matching time points to the intervention group.

**Blood collection for plasma and PBMC isolation**

Blood from participants was collected in three 7.5 mL lithium heparin sampling tubes (Sarsted, 11.608.001) and two 4.9 mL lithium heparin sampling tubes (Sarsted, 04.1939.001) at each time point. 7.5 mL lithium heparin sampling tubes were centrifuged at 3000 RPM for 10 min. Afterwards, plasma was collected and stored at -80 °C until further processing. Additional steps for blood processing are described in the human cohort #4 section.

**Cortisol measurements**

To measure cortisol concentrations in saliva, we employed synthetic swabs specifically designed for cortisol detection (Sarstedt, 51.1534.500). The swabs were centrifuged at 1,000 × g for 5 minutes at 4°C to separate the saliva. The collected saliva samples were then forwarded to the Institute of Clinical Chemistry and Clinical Pharmacology at the University Hospital Bonn, Germany. Cortisol concentrations in the saliva were analyzed by fully automated electrochemiluminescence immunoassays (ECLIA, Elecsys tests) on a cobas e801 analyzer (Roche Diagnostics, Mannheim, Germany) according to the manufacturer's instructions (Roche Diagnostics, 07027150190). The coefficients of variation for intra-assay and inter-assay precision were 2.51% and 3.62%, respectively.

**Targeted and untargeted metabolomic measurements**

**Plasma sample preparation**

Plasma metabolomics samples (50 µL) were mixed with 500 µL of the of ice-cold extraction mixture (metOH/water, 9/1, -20°C, with a cocktail of internal standards), allowing protein precipitation and metabolite extraction, then vortexed and centrifuged (10 min at 15000 g, 4°C). After centrifugation, supernatants were collected and divided in several fractions, and treated following published protocols[17].

Polyamines, biliary acids and small/polar organic acids analyses were performed by LC-MS/MS with a 1290 UHPLC (Ultra-High Performance Liquid Chromatography) (Agilent Technologies) coupled to a QQQ 6470 (Agilent Technologies).

**Regarding polyamines** analyzed by MRM analysis in positive polarity, the gas temperature was set to 350°C with a gas flow of 12 L/min. The capillary voltage was set to 2.5 kV. 10 µL of sample was injected onto a Column Kinetex C18 (150 mm × 2.1 mm particle size 2.6 µm) from Phenomenex, protected by a guard column C18 (5 mm × 2.1 mm) and heated at 40°C by a Pelletier oven. The gradient mobile phase consisted of water with 0.1% of Heptafluorobutyric acid (HFBA, Sigma-Aldrich) (A) and acetonitrile with 0.1% of HFBA (B) freshly made. The flow rate was set to 0.4 mL/min, and gradient as follows: initial condition was 95% phase A and 5% phase B. Molecules were then eluted using a gradient from 5% to 30% phase B over 7 min. The column was washed using 90% mobile phase B for 2.25 min and equilibrated using 5% mobile phase B for 4 min (https://www.ncbi.nlm.nih.gov/pmc/articles/PMC10891084/).

**Regarding biliary acids** analyzed by MRM analysis in negative polarity, gas temperature was set to 310°C with a gas flow of 9 L/min. Capillary voltage was set to 4.5 kV. 10 µl of sample was injected onto a Column Poroshell 120 EC-C8 1200bars (P/N 981758-902, 100 mm × 2.1 mm particle size 1.9 µm) from Agilent technologies, protected by a guard column XDB-C18 (5 mm × 2.1 mm particle size 1.8 µm) and heated at 40°C by a Pelletier oven. Gradient mobile phase consisted of water with 0.2% of formic acid (A) and acetonitrile/isopropanol (1/1; v/v) (B) freshly made. Flow rate was set to 0.5 mL/min, and gradient as follow: initial condition was 70% phase A and 30% phase B, changing to 38% phase B over 2 min. Phases proportion was still over 2 min, then molecules were eluted using a gradient from 38% to 60% phase B over half a minute. Column was washed using 98% mobile phase B for 2 min and equilibrated using 30% mobile phase B for 1.5 min (https://www.ncbi.nlm.nih.gov/pmc/articles/PMC10891084/).

**Regarding the small organic acid ketone bodies and polar metabolites** (so-called SF method) analyzed by MRM analysis in both positive and negative polarity, alternatively, gas temperature was set to 300°C with a gas flow of 12 L/min. Capillary voltage was set to 4 kV in positive and 5 kV in negative polarity mode. 10 µL of sample was injected on a column Zorbax Eclipse XDB- (P/N 981758-902, 100 mm × 2.1 mm particle size 1.8 µm 1200 bars) from Agilent technologies, protected by a guard column XDB-C18 (5 mm × 2.1 mm particle size 1.8 µm) and heated at 50°C by a Pelletier oven. Gradient mobile phase consisted of water with 0.1% of formic acid (A) and 0.1% of formic acid in acetonitrile (B) freshly made. Flow rate was set to 0.7 mL/min, and gradient as follow: initial condition was 80% phase A and 20% phase B for 3 min, changing to 45% phase B over 4 min before rinsing phase and equilibrium before next injection.

Widely targeted by GC-MS/MS was performed on an Agilent 7000C, and pseudo-targeted analysis by UHPLC-HRAM (Ultra-High Performance Liquid Chromatography—High Resolution Accurate Mass) was performed on a U3000 (Dionex)/Orbitrap q-Exactive (Thermo) coupling, previously described in [18,19]. All targeted treated data were merged and cleaned with a dedicated R (version 4.0) package (@Github/Kroemerlab/GRMeta).

**Targeted metabolomic measurements of plasma polyamines**

In addition to the metabolomic workflows described above, plasma samples of cohort #1 (jumpers and controls) were also analyzed for targeted ployamine levels as described in the methods section “Mouse”.

**Cohort #2 – Identification of Biotypes in the Spectrum of Depression (IBIS)**

**Study design**

The study aimed to identify biomarkers for depression and anxiety disorders through a multidimensional approach, including MRI imaging, genotyping, and immunology. These parameters are not included in this manuscript and will be published later. For this manuscript, only the blood samples from these participants were analyzed and compared. It was an observational longitudinal study conducted without therapeutic intervention at the University Hospital Jena in Germany. Participants were recruited between May 2021 and February 2023. A total of 200 participants were included in the study, and blood samples were obtained from 190 of them. 12 participants were later excluded for not meeting the inclusion criteria, including one individual who was diagnosed with schizophrenia during the study. As a result, the final analysis presented in this manuscript is based on data from 178 participants.

**Ethics**

The study was carried out at the Department of Psychiatry and Psychotherapy, University Hospital Jena, Germany and the trial protocol was approved by the Ethics Committee of the University Hospital Jena, Germany (Bachstraße 18, 07743, Jena, ethical approval number: 2021-2158-BO). The study was conducted in accordance with the Declaration of Helsinki and written informed consent was provided by all participants before starting the on-site screening.

**Participants**

Participants were recruited from the Department of Psychiatry and Psychotherapy, the general psychiatric day clinics, and the psychiatric outpatient clinic. Healthy volunteers were recruited through press releases. In the baseline examination, one third of the patients were from the clinically healthy population, one third from the outpatient or general psychiatric day clinics with subclinical or mild symptoms, and one third from inpatient treatment with manifest disorders of varying severity.  Participants were either clinically healthy or had a disorder from the anxious depressive spectrum according International Statistical Classification of Diseases and Related Health Problems, tenth edition (ICD-10), including generalized anxiety disorder, depressive episode, somatization disorder, adjustment disorder, or psychoreactive disorder. They were aged between 18 and 45 years.

Exclusion criteria included harmful use or dependence on psychotropic substances, manifest neurological or rheumatological pre-existing conditions, history of traumatic brain injury with loss of consciousness, untreated chronic internal diseases, non-MRI-compatible metallic implants, intellectual disability, and inability to provide informed consent.

**Randomization, masking and bias minimization**

There was no randomization involved in this observational study. There was no masking in this study. Bias minimization was addressed by recruiting participants from a variety of clinical settings and ensuring a continuum of symptom severity. Data collection was standardized and involved both self-ratings and clinician-administered ratings, using established questionnaires like the BDI and the HAM-D, and semi-structured interviews.

**Procedures**

After providing informed consent, participants underwent anamnesis, diagnostic, and psychometric data collection, partly via psychiatrist-administered ratings and partly through self-ratings. Appointments for these assessments were scheduled either by phone or in person for inpatient participants. Self-rating questionnaires were distributed with appropriate instructions.

For eligible participants in the study, laboratory tests were conducted during a standardized time window between 7:00 and 10:00 AM while fasting. This included routine blood testing (such as blood count, renal, and hepatic function) as well as the isolation of serum at 1,000 × g for 20 minutes. Afterwards, the serum was stored at -80 °C until further processing.

#### Statistical analysis

Differences between groups relative to the main outcome measures were assessed for statistical significance using stepwise linear regression analysis of covariance (ANCOVA) using JMP Pro v16.1.0 (SAS Institute Inc., Cary, NC). Logarithmic transformation was applied to values when not normally distributed. Potential confounding variables were tested systematically for their effects on main outcome measures and included in the model if they significantly improved goodness-of-fit. For a list of confounding variables see **Supplementary Table 5**. Covariates found to significantly affect outcome measures are reported.

### Cohort #3 – Postmortem brains

#### Ethics

All human postmortem sample procedures were approved by the Institutional Review Boards of the University of Mississippi Medical Center, Jackson, Mississippi and the University Hospitals Cleveland Medical Center, Cleveland, Ohio (ethical approval number: UMMC IRB #1999-1002), and are in accordance with the Declaration of Helsinki. Informed consent from the legally defined next of kin was obtained for the collection of tissue, medical records, and retrospective psychiatric interviews.

#### Study design and statistical analysis

Coronal serial sections (14 mm) from fresh frozen postmortem hippocampal blocks were used for these studies from 24 control subjects, 21 subjects with depression, 20 subjects with substance use disorder (SUD) and 18 subjects with comorbid Depr./SUD (**Supplementary Table 6**). Differences between groups relative to the main outcome measures were assessed for statistical significance using stepwise linear regression analysis of covariance (ANCOVA) using JMP Pro v16.1.0 (SAS Institute Inc., Cary, NC). Logarithmic transformation was applied to values when not normally distributed. Potential confounding variables were tested systematically for their effects on main outcome measures and included in the model if they significantly improved goodness-of-fit. For a list of confounding variables see **Supplementary Table 6**. Covariates found to significantly affect outcome measures are reported. Additional details on sample collection and general information can be found in the supplemental materials of Valeri et al.[20].

### Cohort #4 - StressLess Study

#### Study design

The aim of this study was to compare the effect of a 3-week dietary SPD supplementation (through a spermidine-rich wheat germ extract) on depressive symptoms in mildly to moderately depressed patients. It was a monocenter, randomized, double-blind, placebo-controlled study. Participants were recruited between January 2021 and July 2022. To assess compatibility and side effects, the study was initially conducted with healthy participants.

A total of 85 participants were initially included. One participant from the healthy group was excluded due to fainting during blood withdrawal. In the depressed group, five patients were excluded: one due to an Attention-Deficit/Hyperactivity Disorder diagnosis, one because of borderline personality disorder, one due to a sexually transmitted disease, one because of a loss of motivation, and one for starting smoking. Consequently, the final analysis presented in this manuscript is based on data from 79 participants (**Supplementary Table 7**).

Additionally, in this study, sleep data were also collected using questionnaires, sleep diaries, and actigraphy. However, these parameters are not included in this manuscript and will be published later.

#### Ethics

The study was carried out at the Department of Psychiatry and Psychotherapy, University Hospital Bonn, Germany, and the trial protocol was approved by the Ethics Committee of the University Hospital Bonn (Venusberg Campus 1, 53127 Bonn, lfd. Nr. 319/20). Additionally the study was registered with ClinicalTrials.gov (ClinicalTrials identifier: NCT04823806). The study was in accordance with the Declaration of Helsinki and written informed consent was provided by all participants before starting the on-site screening.

#### Participants

Participants were recruited through print and internet advertisements, department websites, social media, and by word of mouth. Local psychiatrists and psychotherapists were also informed about the trial and asked to refer patients for study participation. Evaluation of study eligibility was performed during study enrollment, including a telephone screening and an on-site screening. The criteria for inclusion consisted of meeting the ICD-10 criteria for current mild or moderate depression with a symptom duration of ≥ 2 weeks at screening. Diagnosis was confirmed through psychiatric expert assessments, and the HAM-D 17 was used as an observer rating scale, while the BDI served as a self-rating scale for validation. As per the German S3 guidelines within the HAM-D 17, a score of 9 to 16 indicates mild depression, while a score of 17 to 24 indicates moderate depression. In the case of the BDI, scores ranging from 10 to 19 represent mild depression, and scores from 20 to 29 signify moderate depression. All instruments are well established internationally or in German-speaking countries and have adequate reliability and validity. Exclusion criteria consisted of psychotic disorders, bipolar disorders, eating disorders, major depressive disorder, anxiety disorders, personality disorders, Attention-Deficit/Hyperactivity Disorder, alcohol and substance use disorder, pregnancy or lactation, and regular medication (except for oral contraceptives and thyroid medication). In addition, people who smoked in the 6 months before study inclusion and patients who were treated with an antidepressant or received psychotherapy intervention within 12 months before the study, were also excluded. To minimize the risk of including patients with age-related cognitive impairments, individuals older than 65 years were excluded. The same exclusion criteria were applied to the healthy subjects.

**Dietary supplement**

Participants in the verum group received a spermidine-rich dietary supplement (CelVio® Complex, SPD) extracted from wheat germ. The daily dose of administered plant extract was equivalent to approximately 6 mg spermidine per day. This was administered as three sachets of 2 mg spermidine equivalent each. In addition to spermidine, each sachet also contained approximately 0.6 mg of spermine and 0.3 mg of putrescine. As a comparator condition, the placebo group received spermidine-free rice extract. All study sachets (spermidine and placebo) were identical in all aspects (shape, color, taste, and smell). Study sachets, including placebos, were provided by TLL (The Longevity Labs, Graz, Austria) free of charge. TLL had no influence or insight into the study design, goals, execution, data collection, analysis or interpretation. TLL did not provide financial support for the study.

**Randomization, masking and bias minimization**

Following baseline assessment, participants were randomly assigned to SPD-based supplementation (target intervention) or placebo supplementation (control intervention). A computer-based algorithm was used to generate a block randomization sequence to ensure the balance of the number of patients assigned to target and control intervention. Race and ethnicity data were not tracked in this study as there is no evidence that this might affect our outcomes. Participant allocation was conducted in a 1:1 ratio by a study collaborator who had no involvement in outcome assessments, ensuring blinding. Participants and site staff were blinded to participant group assignment until after the last patient finished participation.

A small team of medical doctors was selected among the study team for the recruitment and monitoring of the participants throughout the study. They saw each participant on a weekly basis and conducted the HAM-D 17 rating at baseline and follow-up appointment within the study arm of depressed participants. This was to establish an environment of comfort and trust for all participants and also to ensure that changes in symptom severity were not overseen. After every appointment, outcome measures were collected and stored in a dedicated database that was separate from the unblinded database.

**Procedures**

Following an initial phone screening, participants provided written informed consent and underwent further screening assessments for eligibility. Within the depressed cohort, these included the BDI, the HAM-D 17, the Resilience Scale[21] and the German version of the WEMWBS[22]. Healthy participants were evaluated by a psychiatrist and classified as "healthy". Additionally, medical history, pre-study medications and demographic information were obtained. In addition to the standardized psychometric assessment a physical examination and a routine blood testing (including blood count, renal and hepatic function) was conducted. Eligible participants were enrolled in the study and randomly assigned to target or control intervention. The intervention period consisted of three weeks of either SPD supplementation or placebo control supplementation. Following a post-intervention visit one week after the intervention period. Enrolled participants were seen on a weekly-basis by the study team at the University Hospital Bonn in Germany. Participants were requested not to eat and not to drink (except for water) anything for at least 12 hours before the weekly appointments. During every appointment eligibility criteria and signs of adverse events were assessed and participants were asked to fill out questionnaires to monitor changes in symptom severity. In addition to the standardized psychometric assessment, a physical examination and blood was taken for either routine blood testing (including complete blood count, renal and hepatic function tests), PBMC isolation or plasma separation for molecular profiling.

**Blood collection for plasma and PBMC isolation**

Blood samples from participants were collected in two 9 mL EDTA (Sarsted, 02.1066.001) and two 4.9 mL lithium heparin sampling tubes (Sarsted, 04.1939.001) each time point. EDTA tubes were centrifuged at 3000 RPM for 10 min. Subsequently plasma was collected and stored at -20 °C until further processing. In parallel, we conducted an autophagy monitoring assay following the procedures outlined in[23]. In one of the two 4.9 mL lithium heparin tubes, CQ (Sigma, C6628) was added at a final concentration of 100 µM, while the other tube received PBS (Thermo Fisher, 14190169) as a control. The lithium heparin tubes were incubated for 1 h in a water bath at 37 °C to allow for the blood to incubate with CQ. After incubation, the lithium heparin tubes as well as the rest of the EDTA blood were carefully loaded on Pancoll solution (PAN Biotech, P04-60500) in Leucosep tubes (Greiner BioOne, 2007473/2026886) and centrifuged at 684 x g for 30 min (brakeless running down). PBMCs were enriched by selecting the interphase of the Pancoll gradient. The PBMCs of the interphase were washed three times with ice-cold PBS. Pellets were stored at -20 °C until they were further processed.

**Cell lysis of PBMCs**

Cell pellets were lysed in an appropriate volume of SDS lysis buffer (50 mM Tris-HCl; Carl Roth, 4855.2; 1% SDS; Carl Roth, 1057.1; pH 8.0), freshly supplemented with 1x cOmplete™

EDTA-free protease inhibitor cocktail (Roche, 4693132001) and 1× PhosSTOP™ phosphatase inhibitor cocktail (Roche, 4906837001) for 20 min on ice. Subsequently, samples were prepared for separation with an SDS-PAGE system by the addition of 4× Laemmli sample buffer (Bio-Rad, 1610747) containing 100 mM DTT (Carl Roth, 6908.1) and heated at 95 °C for 5 min.

**Sample quantification**

See "Sample quantification" in section "Mouse".

**Sample preparation**

Samples were prepared for separation with an SDS-PAGE system by the addition of 4× Laemmli sample buffer (Bio-Rad, 1610747) containing 100 mM DTT (Carl Roth, 6908.1) and heated at 95 °C for 5 min.

**Immunoblotting**

See "Immunoblotting" in the section "Mouse". For the autophagy monitoring assay LC3B (1:1000, Cell Signaling Technologies, #2775) and Hsp70 (1:1000, Cell Signaling Technologies, #4876) were used as primary antibodies.

**Detection of polyamines and other metabolites in clinical samples**

See "Plasma and tissue polyamines" in section "Mouse".

**Appointments with participants**

Weekly appointments differed mainly in the number of questionnaires and participants' questions concerning the study. At the first appointment (T0) baseline assessment and screening assessments for eligibility were conducted which included BDI, HAM-D 17, Resilience Scale, and WEMWBS. The second appointment (T1) focused primarily on tolerability of the supplement and on questions that arose during the first week of participation. In addition, participants filled out the BDI. During the third appointment (T2) participants were asked to fill out the BDI, the Resilience Score, as well as the WEMWBS. On the fourth (T3) BDI was collected by the study team. At the follow-up appointment (T4) baseline assessments were repeated including the HAM-D 17 which was conducted by the same person as the initial assessment. Additionally, participants were offered to get further medical help and support if needed, and more than half of the participants were seen as outpatients at the Department of Psychiatry and Psychotherapy at the University Hospital Bonn, Germany after their participation in the study. Most common reasons for participants to make an appointment as outpatients were psychoeducation for depression, information on antidepressants and relapse prevention.

Cases of non-compliance, protocol deviations, loss to follow-up, and other reasons for participant dropout were always assessed by the study team.

**Statistical analysis**

Participants were randomized in a blinded fashion with 1:1 allocation as described in the section on randomization, masking and bias minimization above.

All datasets were checked for the normality assumption using Shapiro-Wilk tests. In the case of non-normal data distribution, log or square root transformation was performed. The datasets that maintained the violation of the normality assumption even after transformation steps, were analyzed instead using suitable statistical non-parametric tests. For analyses involving multiple groups and repeated measurements, mixed-effects models along with Sidak's multiple comparisons tests ($\alpha = 0.05$) were used. For independent-group comparisons, t-tests or Mann-Whitney tests were conducted, depending on normality. Spearman's rank correlation was employed for correlation analyses. GraphPad Prism (versions 9.0 or 10.0) and SPSS (version 28) were used for statistics and data visualization. Outlier analysis was done by the ROUT method (Q = 1%).

**Cohort #5 – The Early Medication Change (EMC) trial**

**Study design**

All patients were enrolled in the "Randomized clinical trial comparing an early medication change (EMC) with treatment as usual (TAU) in patients with depression – the EMC trial (ClinicalTrials.gov identifier: NCT00974155). A total of 889 depressed patients were enrolled in the study between 2009 and 2014. Details of the study protocol, as well as additional investigations have been described previously[24–26]. The detailed treatment algorithm can be openly accessed by https://trialsjournal.biomedcentral.com/articles/10.1186/1745-6215-11-21. In summary, the EMC trial was a multicenter, randomized, controlled clinical trial, which investigated whether patients with no improvement after 14 days of escitalopram treatment would benefit from an early medication switch (EMC: switch to venlafaxine from day 14, followed by augmentation with lithium if again nonresponsive on day 28) compared with patients treated according to current guidelines (TAU: continue escitalopram for two weeks and switch to venlafaxine on day 28).The main inclusion criteria for the EMC trial were: (1) depression, first episode or recurrent, according to DSM-IV, (2) a HAM-D 17 score of 18 at screening, (3) age 18-65 years and 60 years at the time of the first depressive episode. Main exclusion criteria: (1) bipolar disorder or psychotic depression, (2) benzodiazepines >1.5 mg lorazepam per day, (3) non-native speaker of German, (4) adequate (dose and time) treatments with escitalopram, venlafaxine, or lithium in advance.

**Ethics**

All participants provided written informed consent to participate in the study after a complete and detailed description. All components of the study were approved by the local ethics committee of the Landesärztekammer of Rhineland-Palatinate (study code: 837.166.09 (6671)) and comply with the Declaration of Helsinki in its current version.

**Sample**

The data shown here is is a secondary analysis of 169 depressed patients (Supplementary Table 8) who participated in the EMC trial (n = 889). Blood samples were available at baseline for 560 patients, from which the clearest responders (n = 86) and non-responders (n = 83) were selected. The selection was based on their response to antidepressant medication, measured by the HAM-D 17 after four weeks. In addition, depression severity was assessed at weekly intervals from baseline to week 8. The non-responders showed no improvement in depressive symptoms despite four weeks of antidepressant treatment; in the responders group, depressive symptoms decreased steadily during the course of treatment.

**Study procedures**

At the screening visit, all participants underwent a detailed examination. Diagnoses were verified using the German version of the Mini International Neuropsychiatric Interview (M.I.N.I.)[27] and the Structured Clinical Interview for DSM-IV Axis II Personality Disorders (Münster, 1999)[28]. Sociodemographic and clinical characteristics were assessed by self-report. Depression severity was monitored weekly from baseline to week 8 using the HAM-D 17 by trained and blinded raters. Morning blood samples were taken weekly in fasting patients before medication intake. If necessary, antidepressant premedication was washed out before the baseline visit. From baseline to day 14, antidepressant treatment for all patients was escitalopram (up to 20 mg per day). For the following six weeks, patients were treated according to the study protocol. No changes were made to concomitant medication for general medical conditions.

**References Methods**


1. van der Kooij MA, Jene T, Treccani G, et al. Chronic social stress-induced hyperglycemia in mice couples individual stress susceptibility to impaired spatial memory. *Proc Natl Acad Sci U S A*. 2018;115(43):E10187-E10196. doi:10.1073/pnas.1804412115

2. Hartmann J, Wagner KV, Liebl C, et al. The involvement of FK506-binding protein 51 (FKBP5) in the behavioral and neuroendocrine effects of chronic social defeat stress. *Neuropharmacology*. 2012;62(1):332-339. doi:10.1016/j.neuropharm.2011.07.041

3. Norman KJ, Seiden JA, Klickstein JA, et al. Social stress and escalated drug self-administration in mice I. Alcohol and corticosterone. *Psychopharmacology (Berl)*. 2015;232(6):991-1001. doi:10.1007/s00213-014-3733-9

4. Golden SA, Covington HE, Berton O, Russo SJ. A standardized protocol for repeated social defeat stress in mice. *Nat Protoc*. 2011;6(8):1183-1191. doi:10.1038/nprot.2011.361

5. Newman EL, Covington HE, Suh J, et al. Fighting Females: Neural and Behavioral Consequences of Social Defeat Stress in Female Mice. *Biol Psychiatry*. 2019;86(9):657-668. doi:10.1016/j.biopsych.2019.05.005

6. Lauer J, Zhou M, Ye S, et al. Multi-animal pose estimation, identification and tracking with DeepLabCut. *Nat Methods*. 2022;19(4):496-504. doi:10.1038/s41592-022-01443-0

7. Goodwin NL, Choong JJ, Hwang S, et al. Simple Behavioral Analysis (SimBA) as a platform for explainable machine learning in behavioral neuroscience. *Nat Neurosci*. 2024;27(7):1411-1424. doi:10.1038/s41593-024-01649-9

8. Duque-Wilckens N, Steinman MQ, Busnelli M, et al. Oxytocin Receptors in the Anteromedial Bed Nucleus of the Stria Terminalis Promote Stress-Induced Social Avoidance in Female California Mice. *Biol Psychiatry*. 2018;83(3):203-213. doi:10.1016/j.biopsych.2017.08.024

9. Newman EL, Covington HE, Leonard MZ, Burk K, Miczek KA. Hypoactive Thalamic Crh+ Cells in a Female Mouse Model of Alcohol Drinking After Social Trauma. *Biol Psychiatry*. 2021;90(8):563-574. doi:10.1016/j.biopsych.2021.05.022

10. Hughes CS, Sorensen PH, Morin GB. A Standardized and Reproducible Proteomics Protocol for Bottom-Up Quantitative Analysis of Protein Samples Using SP3 and Mass Spectrometry. *Methods Mol Biol*. 2019;1959:65-87. doi:10.1007/978-1-4939-9164-8_5

11. Bian Y, Zheng R, Bayer FP, et al. Robust, reproducible and quantitative analysis of thousands of proteomes by micro-flow LC-MS/MS. *Nat Commun*. 2020;11(1):157. doi:10.1038/s41467-019-13973-x

12. Cox J, Hein MY, Luber CA, Paron I, Nagaraj N, Mann M. Accurate proteome-wide label-free quantification by delayed normalization and maximal peptide ratio extraction, termed MaxLFQ. *Mol Cell Proteomics*. 2014;13(9):2513-2526. doi:10.1074/mcp.M113.031591

13. Tyanova S, Temu T, Sinitcyn P, et al. The Perseus computational platform for comprehensive analysis of (prote)omics data. *Nat Methods*. 2016;13(9):731-740. doi:10.1038/nmeth.3901

14. Goedhart J, Luijsterburg MS. VolcaNoseR is a web app for creating, exploring, labeling and sharing volcano plots. *Sci Rep*. 2020;10(1):20560. doi:10.1038/s41598-020-76603-3

15. Magnes C, Fauland A, Gander E, et al. Polyamines in biological samples: rapid and robust quantification by solid-phase extraction online-coupled to liquid chromatography-tandem mass spectrometry. *J Chromatogr A*. 2014;1331(100):44-51. doi:10.1016/j.chroma.2013.12.061

16. Costa-Machado LF, Garcia-Dominguez E, McIntyre RL, et al. Peripheral modulation of antidepressant targets MAO-B and GABAAR by harmol induces mitohormesis and delays aging in preclinical models. *Nat Commun*. 2023;14(1):2779. doi:10.1038/s41467-023-38410-y

17. Grajeda-Iglesias C, Durand S, Daillère R, et al. Oral administration of Akkermansia muciniphila elevates systemic antiaging and anticancer metabolites. *Aging (Albany NY)*. 2021;13(5):6375-6405. doi:10.18632/aging.202739

18. Abdellatif M, Trummer-Herbst V, Koser F, et al. Nicotinamide for the treatment of heart failure with preserved ejection fraction. *Sci Transl Med*. 2021;13(580):eabd7064. doi:10.1126/scitranslmed.abd7064

19. Durand S, Grajeda-Iglesias C, Aprahamian F, Nirmalathasan N, Kepp O, Kroemer G. The intracellular metabolome of starving cells. *Methods Cell Biol*. 2021;164:137-156. doi:10.1016/bs.mcb.2021.04.001

20. Valeri J, Stiplosek C, O'Donovan SM, et al. Extracellular matrix abnormalities in the hippocampus of subjects with substance use disorder. *Transl Psychiatry*. 2024;14(1):115. doi:10.1038/s41398-024-02833-y

21. Wagnild GM, Young HM. Development and psychometric evaluation of the Resilience Scale. *J Nurs Meas*. 1993;1(2):165-178.

22. Tennant R, Hiller L, Fishwick R, et al. The Warwick-Edinburgh Mental Well-being Scale (WEMWBS): development and UK validation. *Health Qual Life Outcomes*. 2007;5:63. doi:10.1186/1477-7525-5-63

23. Bensalem J, Hattersley KJ, Hein LK, et al. Measurement of autophagic flux in humans: an optimized method for blood samples. *Autophagy*. 2021;17(10):3238-3255. doi:10.1080/15548627.2020.1846302

24. Tadić A, Gorbulev S, Dahmen N, et al. Rationale and design of the randomised clinical trial comparing early medication change (EMC) strategy with treatment as usual (TAU) in patients with major depressive disorder--the EMC trial. *Trials*. 2010;11:21. doi:10.1186/1745-6215-11-21

25. Tadić A, Wachtlin D, Berger M, et al. Randomized controlled study of early medication change for non-improvers to antidepressant therapy in major depression--The EMC trial. *Eur Neuropsychopharmacol*. 2016;26(4):705-716. doi:10.1016/j.euroneuro.2016.02.003

26. Engelmann J, Zillich L, Frank J, et al. Epigenetic signatures in antidepressant treatment response: a methylome-wide association study in the EMC trial. *Transl Psychiatry*. 2022;12(1):268. doi:10.1038/s41398-022-02032-7

27. Sheehan DV, Lecrubier Y, Sheehan KH, et al. The Mini-International Neuropsychiatric Interview (M.I.N.I.): the development and validation of a structured diagnostic psychiatric interview for DSM-IV and ICD-10. *J Clin Psychiatry*. 1998;59 Suppl 20:22-33;quiz 34-57.

28. Münster RD. Wittchen, H.-U., Zaudig, M. & Fydrich, T. (1997). SKID Strukturiertes Klinisches Interview für DSM-IV. Achse I und II. Göttingen: Hogrefe, DM 158,-. Hiller, W., Zaudig, M. & Mombour, W. (1997). IDCL Internationale Diagnosen Checklisten für DSM-

IV und ICD-10. Göttingen: Hogrefe, DM 198,- bzw. DM 239,-. *Zeitschrift für Klinische Psychologie und Psychotherapie*. 1999;28(1):68-70. doi:10.1026//0084-5345.28.1.68
